# Cortical network structure mediates response to stimulation: an optogenetic study in non-human primates

**DOI:** 10.1101/2021.05.17.444526

**Authors:** Julien Bloch, Alexander Greaves-Tunnell, Eric Shea-Brown, Zaid Harchaoui, Ali Shojaie, Azadeh Yazdan-Shahmorad

## Abstract

As aberrant network-level functional connectivity underlies a variety of neural disorders, the ability to induce targeted functional reorganization would be a profound development towards therapies for neural disorders. Brain stimulation has been shown to alter large-scale network-wide functional connectivity, but the mapping from stimulation to the modification is unclear. Here, we leverage advances in neural interfaces, interpretable machine learning, and graph theory to arrive at a model which accurately predicts stimulation-induced network-wide functional reorganization. The model jointly considers the stimulation protocol and the cortical network structure, departing from the standard approach which only considers the stimulation protocol. We validate our approach in the primary sensorimotor cortex of non-human primates using paired optogenetic stimulation through a large-scale optogenetic interface. We observe that the stimulation protocol only predicts a small portion of the induced functional connectivity changes while the network structure predicts much more, indicating that the underlying network is the primary mediator of the response to stimulation. We extract the relationships linking the stimulation and network characteristics to the functional connectivity changes and observe that the mappings diverge over frequency bands and successive stimulations. Finally, we uncover shared processes governing real-time and longer-term effects of stimulation. Our framework represents a paradigm shift for targeted neural stimulation and can be used to interrogate, improve, and develop stimulation-based interventions for neural disorders.

**Teaser:** Brain stimulation rewires the brain, but the pre-existing network structure of the brain controls the rewiring.

## Introduction

From schizophrenia to epilepsy, aberrant network-level functional connectivity underlies a variety of neural disorders (*1*–*3*). As such, the ability to modify network-level functional connectivity would be a profound development towards therapies for neural disorders. Brain stimulation has demonstrated promise as a means for modifying large-scale functional connectivity (*4*–*6*), but the mapping from stimulation to the resulting modification remains unclear.

The prevalent approach for inducing targeted connectivity change is neural stimulation informed by Spike-Timing Dependent Plasticity (STDP). STDP is a phenomenon in which synaptic strengths are modified as a function of the delay between the firing times of the pre- and post-synaptic neurons (*7*–*9*). While STDP was initially observed in monosynaptic connectivity strength between isolated neuron pairs *in vitro* (*7*–*9*), its application was eventually translated to larger-scale, less isolated connections in vivo (*10*–*16*). In response to the more challenging in vivo setting, this translation coincided with stimulation of a larger set of neurons instead of monosynaptic connections of isolated neuron pairs and the measurement of functional connectivity changes (FCC) instead of direct synaptic strength changes (*10*–*16*). In line with the original in *vitro* studies, these experiments hypothesized that pairwise stimulation would drive FCC only between the stimulation targets. Some reported successful induction of targeted FCC (*13*–*16*) while others reported inconsistent results across stimulation targets (*10*, *11*). Notably, these studies also reported a novel finding: stimulation-induced FCC were found to extend beyond the stimulation sites to other recorded regions (*10*, *11*, *14*–*16*). Follow up studies investigated the extent of this phenomenon in primates (*12*, *17*, *18*) and found that paired stimulation reliably results in FCC extending over a large-scale surrounding network. These results highlight the need for an updated framework which accurately predicts network-wide stimulation-induced FCC *in vivo*.

Recent research indicates that network-level functional connectivity is not only affected by stimulation but also is involved in shaping the response to stimulation. For example, underlying functional connectivity has been shown to shape stimulation-induced neural activity propagation (*18*–*22*) as well as influence the therapeutic outcome of neural stimulation interventions for Parkinson’s disease (*23*, *24*) and depression (*25*). Early results have also indicated that network-level FCC induced by stimulation are correlated to baseline functional connectivity (*11*, *17*, *26*). Despite the demonstrated relevance of the underlying network for shaping the response to stimulation, the question of how to effectively parse and analyze a brain network to inform stimulation remains unanswered. The answer to this question could be used to arrive at an updated framework for network-level stimulation induced FCC.

Here, we leverage advances in neural interfaces and computational modeling to develop a model which jointly considers characteristics of the stimulation protocol and the underlying network-level functional connectivity for prediction of network-wide stimulation-induced FCC. We use optogenetics, a stimulation technology in which neurons are rendered light-sensitive by viral-mediated expression of opsins and thus able to be activated by incident light (*27*), to activate target neural regions while simultaneously recording without electrical artifact. We measure FCC at the network level by recording neural activity via a micro-electrocorticography (μECoG) array covering ~ 1cm^2^ of the primary sensorimotor cortex. We then parse underlying network-level functional connectivity with summarizing metrics borrowed from and inspired by graph theory. Finally, we employ a nonparametric hierarchical additive model (*28*) to predict network-level FCC while allowing maximal interpretability of identified feature mappings. We develop and test the model in the sensorimotor cortex of two non-human primates (NHPs) to ensure maximal clinical translation.

We interrogated our model over multiple contexts to identify the relationship between stimulation and FCC. Characteristics of the stimulation protocol alone poorly predicted stimulation induced FCC, whereas the characteristics of the underlying network-level functional connectivity contained information which yielded accurate prediction of the FCC. This trend was true both when protocol and network features were analyzed as groups and when features were analyzed individually. The mappings from input features to FCC were similar between the stimulated-state and resting-state changes. Continuous stimulation modified the mapping from input features to resulting FCC. Our methods represent a promising new framework for parsing network-level FCC to predict the effects of stimulation, while our model and insights can be leveraged towards novel therapies of neural disorders.

## Results

### Large-scale stimulation and recording of non-human primate sensorimotor cortex

We performed a series of experimental sessions consisting of stimulating and recording rhesus macaque cortex. To stimulate neurons without causing a recording artifact we used optogenetic stimulation. We chose to express the C1V1 opsin as it is a red-shifted excitatory opsin and thus allows for greater light penetration and subsequent neural activation (*29*). To record over a large-scale (~ 1cm^2^) area of the cortex while allowing light to pass through the recording hardware, we used a transparent μECoG array which recorded local-field potentials (LFP) from the surface of the primary sensorimotor cortex (*30*).

For each experimental session we randomly chose two locations corresponding to two electrodes of the μECoG array to be stimulated. The stimulation consisted of alternating 5 ms laser pulses delivered to the stimulation sites, temporally offset by a session-specific delay (Fig. 1) and repeated every 200 ms for 10 minutes. The session-specific delays were randomly chosen at the beginning of each experimental session to be 10 ms, 30 ms, or 100 ms. These values were chosen because 10 ms and 30 ms have been shown to be in the effective range for connectivity change while 100 ms is outside the effective range according to STDP (*9*, *13*). Locations of stimulation, the delay between the stimulation of the first and second sites, and the location and orientation of the μECoG array placement varied between experimental sessions but were consistent over individual sessions.

**Figure 1:**
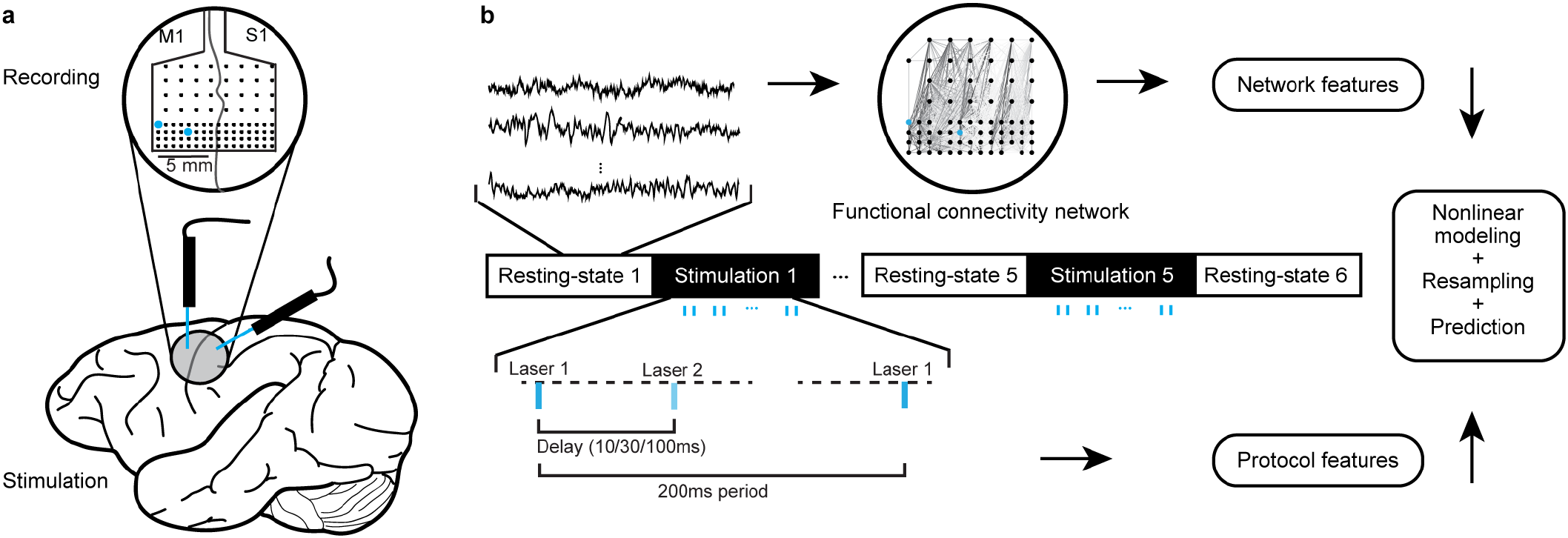
Schematic of the experimental setup and basic modeling pipeline. **a**, Depiction of the transparent 96-electrode μECoG array which we used to record from macaque primary sensorimotor cortex, and the approximate location of the array on the cortex. Optogenetic stimulation was applied via laser light. **b**, We recorded traces of neural LFP during the resting-state and stimulation blocks making up our experimental sessions. During the stimulation blocks, stimulation was applied in a paired pulse protocol whereby the stimulation at the second site followed the first by a session-specific delay. Network coherences were constructed for each individual recording block by calculating pairwise LFP coherences over the network. Network coherence from resting-state recordings were parameterized to generate a set of network features. Details of the protocol was parameterized to generate a set of protocol features. The sets of network and protocol features were fed to a nonparametric model to predict network-wide FCC.

The set of experimental sessions spanned multiple days and two rhesus macaques. We recorded neural activity during 10-minute stimulation periods, which we refer to as “stimulation blocks,” as well as during 5-minute periods before and after each stimulation period, which we refer to as “resting-state blocks.” Each experimental session consisted of alternating resting-state blocks and stimulation blocks, with 6 total resting-state blocks and 5 total stimulation blocks.

From the neural data recordings we calculated the FCC for each pairwise connection between recorded electrodes of an experimental session. We quantified FCC with the coherence metric for the Theta (4-7 Hz), Beta (12-30 Hz), Gamma (30-70 Hz), and High-gamma (70-200 Hz) bands.

As the effects of stimulation in both the stimulated-state and in the resting-state are relevant for neural disorders therapies (*31*, *32*), we estimate and model the FCC in both states. We calculated the change from a resting-state block to the following stimulation block which we refer to as SS-FCC, and calculated the change from a resting-state block to the following resting-state block which we refer to as RS-FCC.

### Stimulation delay is a poor predictor of resting-state functional connectivity change between stimulation sites

Studies have shown that change in anatomical connectivity between two monosynaptically connected neurons is a function of activation times, or stimulation delay, between the neurons (*7*–*9*), a phenomenon termed STDP. As STDP relates delay to structural changes in connectivity but not real-time modulation of neural activity, we evaluated how well stimulation delay could predict RS-FCC between the sites of stimulation.

The standard function relating stimulation delay to corresponding connectivity change according to the STDP hypothesis is an exponential function (*33*, *34*). However, here we recognize that our experimental setting bears little resemblance to the controlled monosynaptic studies yielding such a curve. As such, we did not enforce an exponential delay function but rather tested whether varying the delay had an effect on RS-FCC by fitting a linear model with delay as categorical input and RS-FCC as continuous output (Fig. 2a). Since STDP is a process governing pairwise connectivity change, we modeled only RS-FCC calculated between stimulation sites, and did not model RS-FCC nor connectivity between unstimulated sites. We observed that delay was a significant predictor of RS-FCC in Theta and Gamma bands, but not for Beta or High-gamma (*p*: Theta 0.016, Beta 0.125, Gamma 0.002, High-gamma 0.239; one-sided F-test with 165 observations, 2 degrees of freedom). For all frequencies we observed that delay had low explanatory power (*R*^2^: Theta 0.050, Beta 0.025, Gamma 0.075, High-gamma 0.018). These results indicate that, although a naive extrapolation of the STDP curve to our large-scale setting would hint otherwise, delay between paired stimulation of two neural sites minimally controls RS-FCC.

**Figure 2:**
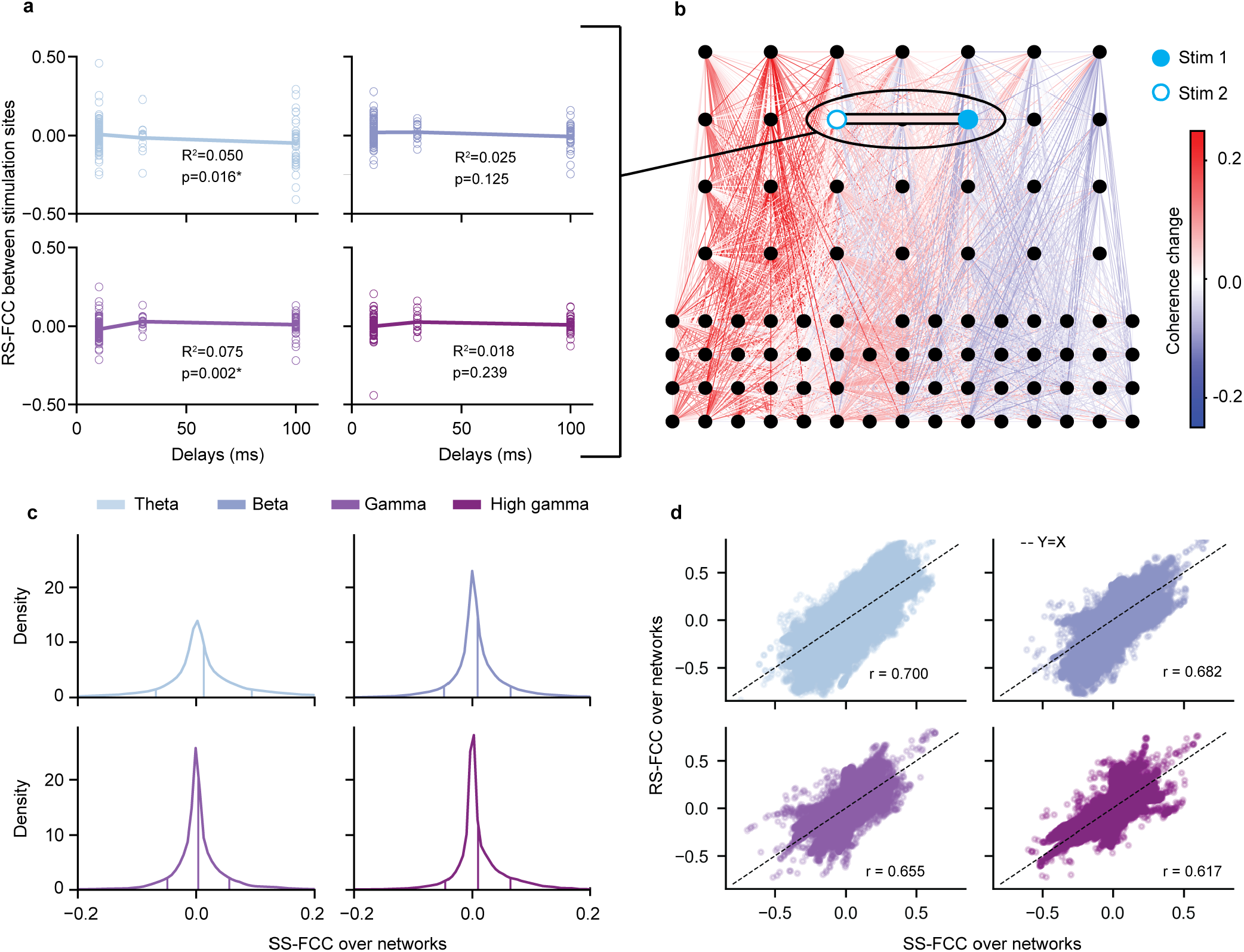
**a**, Delay-based prediction of RS-FCC between stimulation sites has poor explanatory power, and does not achieve statistical significance in the Beta and High-gamma. **b**, Visualization of the RS-FCC of a single example session over the μECoG array. Black circles correspond to electrodes while the edges indicate the change in coherence relative to the preceding baseline period. Optogenetic stimulation sites are indicated by light blue circles. The bolded line connecting the stimulation sites indicates the coherence change between the two sites. **c**, Distribution of the SS-FCC in each frequency band over all sessions. Internal lines indicate the mean, and one standard deviation below and above the mean. **d**, Scatter plot of SS-FCC versus RS-FCC showing high correlation.

### Pairwise stimulation drives functional connectivity changes in the stimulated-state and resting-state across the entire recorded network

In order to determine whether our stimulation induced FCC only between the stimulation sites or over a broader area of the recorded network, we examined the network-wide FCC. We plot an example network of FCC over our recording array in Fig. 2b. In this example, we see that the connectivity change between stimulation sites (bolded) is simply one change among many in the recorded network. To quantify the network-wide FCC over all experiments we calculated the FCC distributions. We observed that stimulation generally increased network-level functional connectivity, as SS-FCC had positive mean over all frequency bands. We additionally compared the SS-FCC of our stimulation experiments to a set of control experiments in which no stimulation was applied, and found that stimulation resulted in significantly higher SS-FCC (*p* < 0.0001 in all frequency bands; Welch’s unequal variances t-test) (Fig. S1.)

We then compared the SS-FCC and RS-FCC distributions. We found SS-FCC was highly correlated to RS-FCC, as the Pearson’s correlation coefficient was above 0.6 in all frequency bands (Fig. 2d). The means of RS-FCC distributions were less offset from 0 than the means of SS-FCC distributions of the same frequency bands, and variances were similar between RS-FCC and SS-FCC distributions of the same frequency bands. These widespread FCC raise the question of explaining and predicting the stimulation-induced FCC across the entirety of the recorded network.

### Nonlinear modeling to predict stimulation-induced network-wide functional connectivity changes

Motivated by the observation of network-wide FCC, we pursued an accurate and interpretable model to predict FCC over the entire recorded network. While linear modeling is the standard method for statistically rigorous interpretable models, the assumption of linearity is scientifically unreasonable for statistical relationships among complex experimental measurements such as neural data (*35*). We relax this assumption through the use of a nonparametric hierarchical additive model which can identify complex nonlinear input-output relationships (*28*). As this model retains the structure of summation over individual features familiar from linear modeling, it permits feature-wise investigation of the results (Fig. 3c-f). In addition, careful penalization of the estimation objective (Eq. 18) ensures that the model retains important theoretical guarantees and is encouraged to learn simple, smooth components (*28*). In the following sections we use such nonparametric hierarchical models to arrive at our results. These models outperformed linear models uniformly across frequency bands and prediction tasks (Fig. S2).

**Figure 3:**
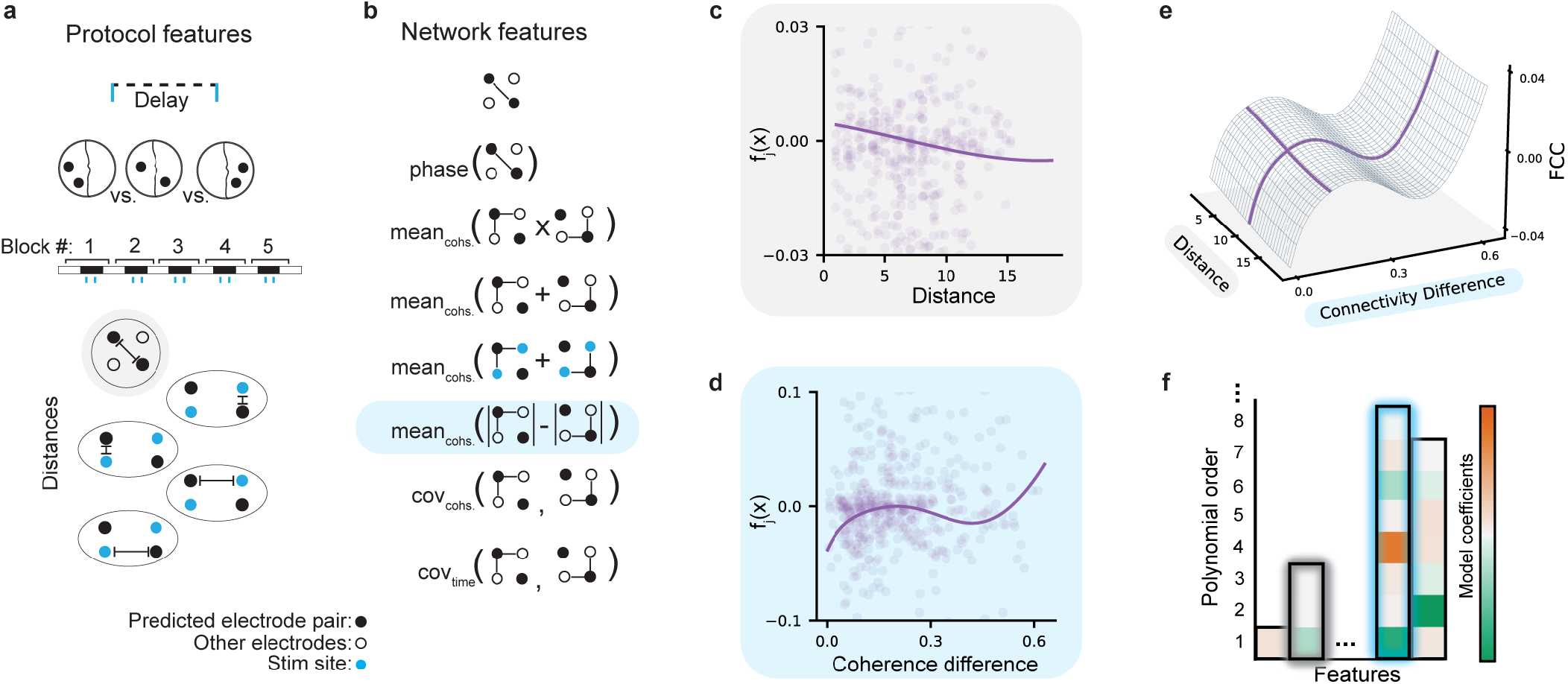
Feature generation and predictive modeling. **a**, Visualization of the protocol features. From top to bottom, the features are: stimulation delay, anatomical region, block number, distance, stim 1 distance to closer, stim 2 distance to closer, stim 1 distance to further, stim 2 distance to further. **b**, Visualization of the network characteristics which the network features quantify. From top to bottom, the visualizations correspond to: initial coherence, phase, length 2 path strength, coherence with network, coherence with stim sites, coherence difference, electrode covariance, and time covariance. **c-d** The nonparametric model can identify feature mappings of a variety of complexities, such as (**c**) near-linear fits such as that found for the distance feature, or (**d**) higher-order polynomials such as that found for the connectivity difference feature. *f_j_*(*x*) indicates the contribution of a feature to the predicted FCC. Throughout the figure, the gray shadowing indicates the distance feature, while the light blue shadowing indicates the connectivity difference feature. **e**, An example of how the feature mappings are additively combined. **f**, Individual feature mappings are represented in a basis of increasingly complex nonlinear functions. The order of this representation is selected automatically by a penalized estimation procedure, with the y-axis of this figure indicating the polynomial order. The order selection for the distance feature and coherence difference feature are shadowed in gray and light blue, respectively.

### The stimulation protocol alone is a poor predictor of network-wide functional connectivity changes

To test the degree to which the factors of the stimulation protocol controlled the network-wide FCC, we constructed models to predict SS-FCC and RS-FCC from the stimulation protocol. We constructed a set of 8 features consisting of: stimulation delay, a categorical identifier of the time ordering of the stimulation block, the distance between a given electrode pair, the distances from the sites of stimulation to the closer or further electrode of an electrode pair, anatomical region of electrodes (Fig. 3a) (see Methods for more details of how they are calculated). Throughout the paper we refer to this set of predictive features as “protocol features.” We also include a categorical animal subject indicator in the model in order to capture differences in mean between the two subjects. With this feature set we trained models with output of pairwise FCC between all recording electrodes of an experiment, over each block of all experiments. We fit individual models to predict SS-FCC or RS-FCC over individual frequency bands. We held out 30% of the data to evaluate the predictive performance of the model on unseen data.

We observed that the protocol-based model of FCC had low predictive power for both SS-FCC (*R*^2^: Theta 0.045, Beta 0.026, Gamma 0.057, High-gamma 0.078) and RS-FCC (*R*^2^: Theta 0.028, Beta 0.014, Gamma 0.020, High-gamma 0.014). These results indicate that while factors related to the protocol alone explain some of the variation in the observed coherence change, they are insufficient to generate practically relevant predictions.

### Network structure determines stimulated-state and resting-state functional connectivity changes

Our observations that stimulation induces a change in connectivity over an entire network, in tandem with the poor predictive power of the protocol-based model, lead us to hypothesize that the effects of stimulation may be mediated by the existing cortical network. We tested this hypothesis by constructing a set of 8 features extracted from the baseline functional connectivity network. These features are: initial coherence, which is the pairwise coherence between an electrode pair; phase, which is the absolute value of difference between electrode spectral phases; length 2 path strength, which is the average strength of a connection between an electrode pair passing through another recording site; coherence with network, which is the average coherence between a given electrode pair and the rest of the network; coherence with stim sites, which is the average coherence between a given electrode pair and the stimulation sites; coherence difference, which is the average absolute difference in connection strength between each element of the electrode pair and the recorded network; electrode covariance, which is the average covariance of an electrode pair’s coherence to the other electrodes in the network over time; and time covariance, which is the average covariance across time series between a given electrode pair’s coherence and the others in the network (Fig. 3b) (see Methods for details of how these are calculated). These features are primarily derived from graph theoretic measures (*36*–*39*) which have been used with success in past theoretical neuroscience analyses (*40*–*43*). Throughout this paper we refer to this set of features as “network features.”

We observed that the model predictions on the test set was dramatically improved compared to the protocol-only model for each frequency band for both SS-FCC (*R*^2^: Theta 0.240, Beta 0.130, Gamma 0.221, High-gamma 0.323) (Fig. 4) and RS-FCC (*R*^2^: Theta 0.218, Beta 0.087, Gamma 0.138, High-gamma 0.218) (Fig. 6a). This improvement in performance across all frequency bands and for both stimulated-state and resting-state contexts suggests that properties of the local network at baseline may be highly informative with respect to prediction of stimulation-induced changes.

**Figure 4:**
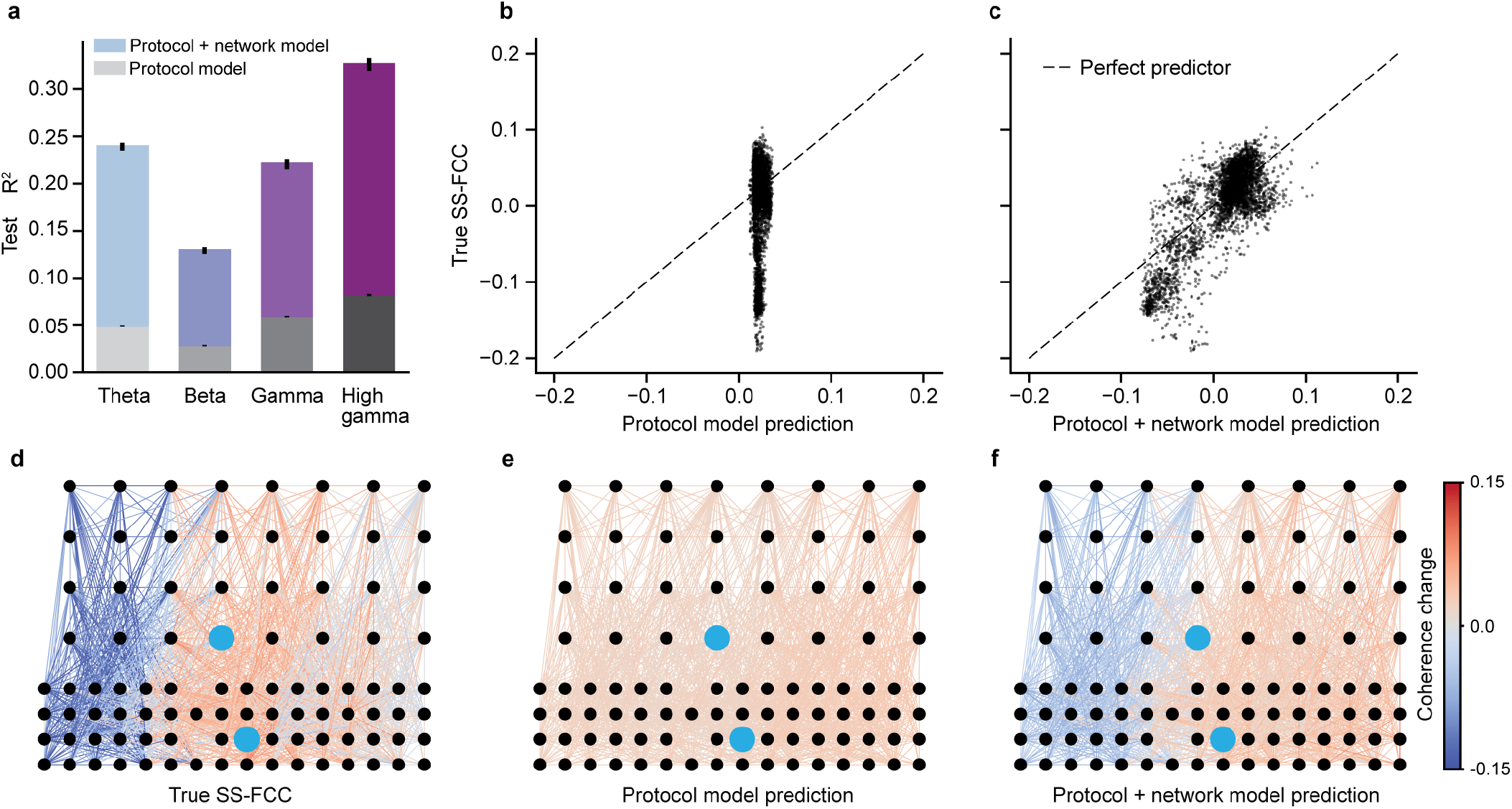
A model trained to predict stimulation-induced FCC over a cortical network based on protocol features alone is outperformed by one which incorporates information about existing network connectivity. Results in this figure correspond to prediction quality for SS-FCC; results are similar but slightly worse for RS-FCC. **a**, Bar graph indicating r-squared accuracy on held-out data. Grayscale bars indicate accuracy of models trained on only protocol features and colored bars indicate accuracy of models trained on both protocol and network features. Error bars indicate standard deviation. **b-f**, Results of prediction of a representative experimental session from the protocol versus the protocol-and-network models. **b**, Scatter plot of the protocol model’s predictions of network SS-FCC versus the actual changes observed during that experimental session. **c**, Scatter plot of the protocol-and-network model’s predictions of SS-FCC versus the actual changes observed during that experimental session. **d**, Example SS-FCC over a cortical network, visualized over the μECoG array used for recording. Black circles indicate electrodes of the array, while lines indicate coherence change between electrode pairs. The observed coherence changes correspond to the y-axis of **b** and **c**. **e**, The protocol model’s SS-FCC prediction, corresponding to the x-axis of **b**. **f**, The protocol-and-network model’s SS-FCC prediction, corresponding to the x-axis of **c**.

### Individual network features are more important than individual stimulation protocol features for predicting functional connectivity changes

We quantified the relative importance of each feature through the decrease in accuracy of prediction on the test set when that feature was excluded from the data. The statistical uncertainty associated with the importance was quantified through a resampling procedure (*44*) (see Methods for details).

Using this procedure, we show the importances of each feature per frequency band in Fig. 5a for SS-FCC and in (Fig. S3) for RS-FCC. Four of the five most important features for SS-FCC prediction and each of the five most important features for RS-FCC prediction were network features. Features appearing in the top five of both contexts were time covariance, coherence difference, and initial coherence, which were all network features. Delay and time-ordered block number were the two most important of the protocol features for both SS-FCC and RS-FCC prediction.

**Figure 5:**
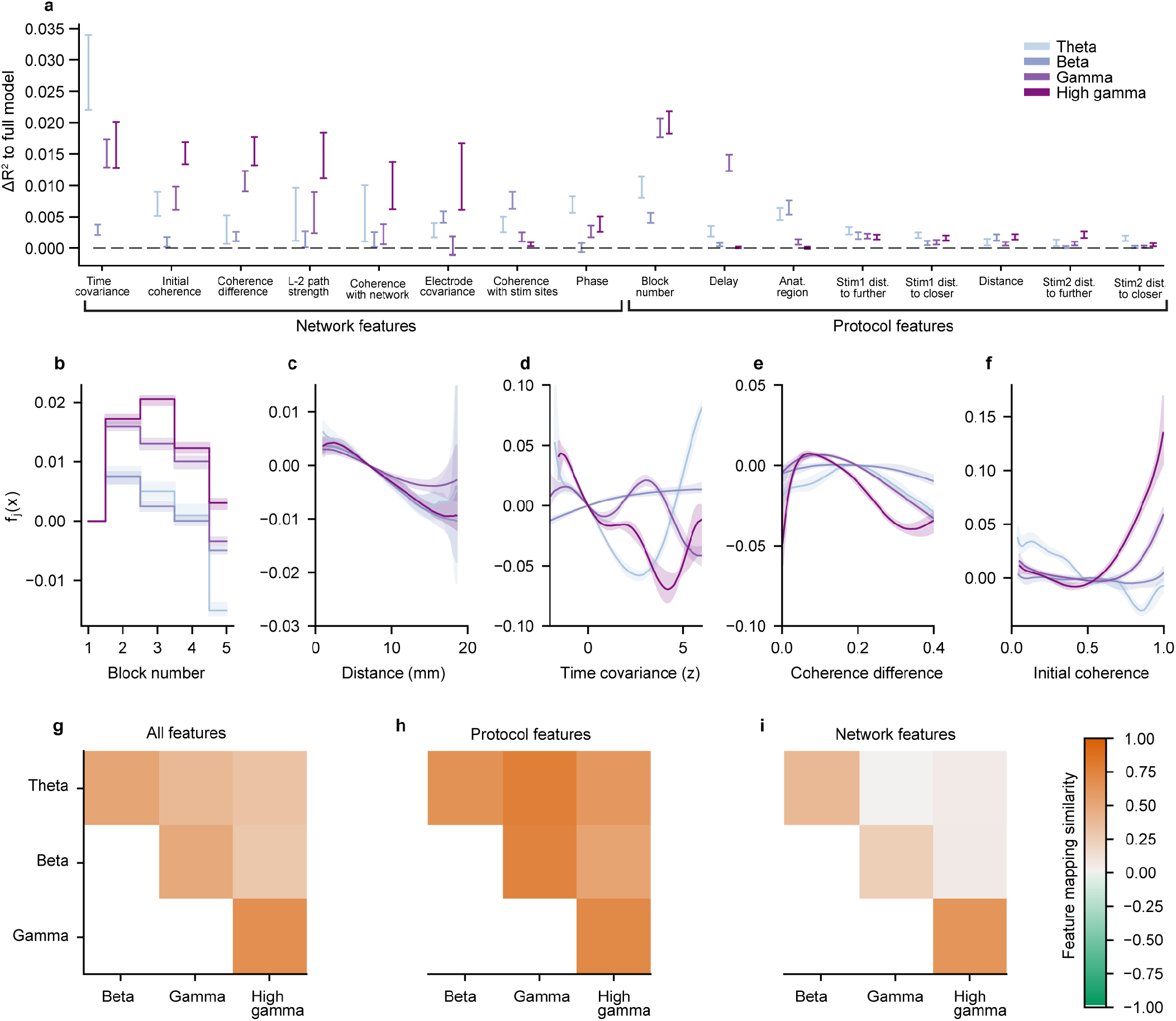
Feature-wise analysis of the nonlinear additive model. Results in this figure correspond to the SS-FCC model; results were similar for the RS-FCC model. **a**, Feature importances as measured by the decrease in model accuracy when the feature was omitted. Features were partitioned into the protocol and network categories, then sorted by average importance over all frequency bands. Line ranges indicate 95% confidence intervals. **b-f**, Feature mappings for the **b** time-ordered block number, **c** electrode pair distance, **d** time covariance (z-scored), **e** coherence difference, and **f** initial coherence features. *f_j_*(*x*) indicates the contribution of a feature to the predicted FCC. Shading indicates 95% confidence intervals. **g-i**, Average cosine similarity of the component functions between each pair of frequency bands, over **g** all features, **h** protocol features, and **i** graph features.

### Feature mappings display diverging trends across frequency bands

We decomposed the nonlinear model into feature mappings representing the estimated mapping between each individual feature and the change in coherence. The statistical uncertainty associated with the estimated nonlinear mapping from the feature to the response was quantified through a resampling procedure (*44*) and representative feature mappings were obtained through point-wise interpolation of the resampled feature mappings (see Methods for details).

We plot the SS-FCC feature mappings of some important features as well as others of interest in Fig. 5b-f. We observed that feature mappings were generally more complicated for network features than for protocol features. For example, the time covariance feature varies from a near-linear increase in the Beta band to a high-order polynomial in the High gamma band, while the distance feature mapping exhibits a nearly linear decrease in the high-confidence regions of all frequency band models (Fig. 5c). We also observed that in the higher frequency bands, electrodes which had a higher initial coherence tended to strongly increase their coherence when stimulated (Fig. 5f). Additionally, we found a small effect of increasing SS-FCC for the first two or three temporal blocks depending on frequency band, followed by decreasing SS-FCC in later blocks (Fig. 5b). We evaluate the temporal characteristics of the FCC in more detail in a later section.

We summarized the similarity of all features, protocol features, and network features by calculating the cosine similarity of the feature mappings (Fig. 5g-i). Generally, we observed that the mappings from protocol features to SS-FCC were fairly consistent across frequency bands, while the mappings from network features to SS-FCC varied sharply between frequency bands. Only the network feature mappings of Gamma and High-gamma were similar.

### Feature mappings are similar for stimulated-state and resting-state functional connectivity changes

We calculated the similarity of feature mappings between SS-FCC and RS-FCC and observed that they exhibited high similarity in all frequency bands (Fig. 6b). Notably, the groups with the highest similarity between SS-FCC and RS-FCC feature mappings were the network features in the Theta and High-gamma bands. We plot representative SS-FCC and RS-FCC feature mappings of five features from the Theta and High-gamma models in Fig. 6c-d.

**Figure 6:**
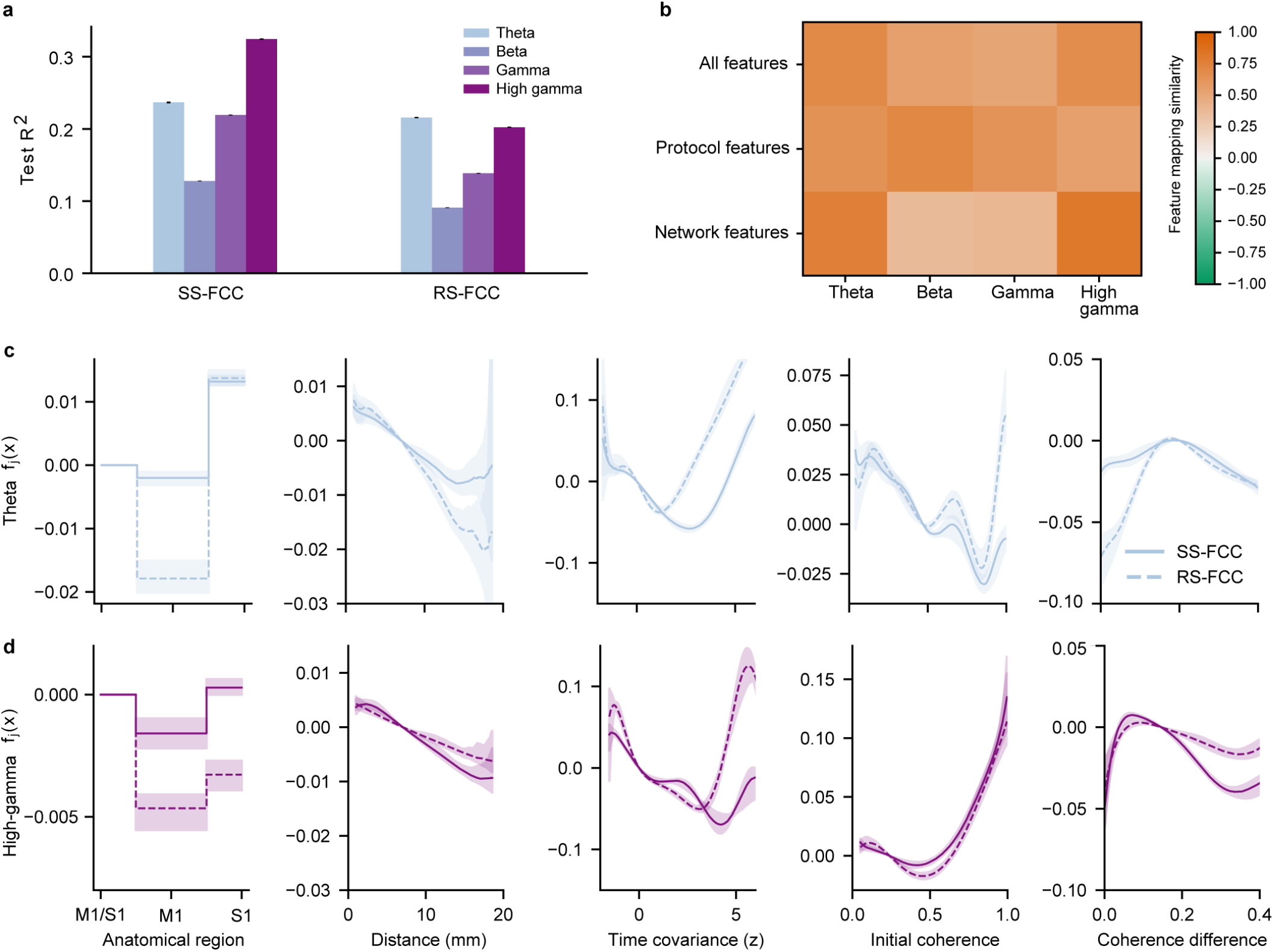
**a**, Comparison of the predictive accuracy for nonlinear additive models of SS-FCC prediction versus RS-FCC prediction. Error bars indicate standard deviation. **b**, Heatmap of average cosine similarity between feature mappings of SS-FCC and RS-FCC over frequencies. **c-d**, Comparison of feature mappings for SS-FCC and RS-FCC models for the **c** Theta band and **d** High-gamma band. The features from left to right are anatomical region of electrodes, electrode pair distance, time covariance (z-scored), initial coherence, and coherence difference. Shading indicates 95% confidence intervals.

### Repeated stimulation modifies mappings from features to functional connectivity changes

The modeling approach employed thus far constrained the learned feature mappings to be constant across all temporal blocks of an experimental session. The influence of the block’s time location was represented by a learned global shift in the FCC between all electrode pairs measured in that block. We hypothesized that successive stimulation blocks within an experiment had a more complex effect on FCC than a simple average shift, and rather modified the actual feature mappings over the time course of stimulation. We tested this hypothesis by allowing the nonlinear relationship between each feature and the response to vary across the five stimulation blocks by adding an interaction term between the block number feature and all other features. Rather than averaging over potential heterogeneity in the network dynamics resulting from the accumulated effects of stimulation, this approach allowed for the investigation of a time-course of smoothly varying predictors of coherence changes.

In line with the hypothesis that repeated stimulation indeed produces such heterogeneity across blocks within a session, we found a substantial improvement in prediction quality with the time-varying model for both SS-FCC (*R*^2^ Theta 0.426, Beta 0.247, Gamma 0.415, High-gamma 0.527) (Fig. 7a) and RS-FCC (*R*^2^ Theta 0.410, Beta 0.251, Gamma 0.359, High-gamma 0.436) over the static model. We also calculated the cosine similarities of feature mappings for each block of the time-varying SS-FCC model and found that successive stimulation results in the mappings diverging from each other, as their similarity decreased over subsequent blocks (Fig. 7d). These results indicate that the relationship between features and connectivity change evolved dynamically over the course of repeated stimulation.

**Figure 7:**
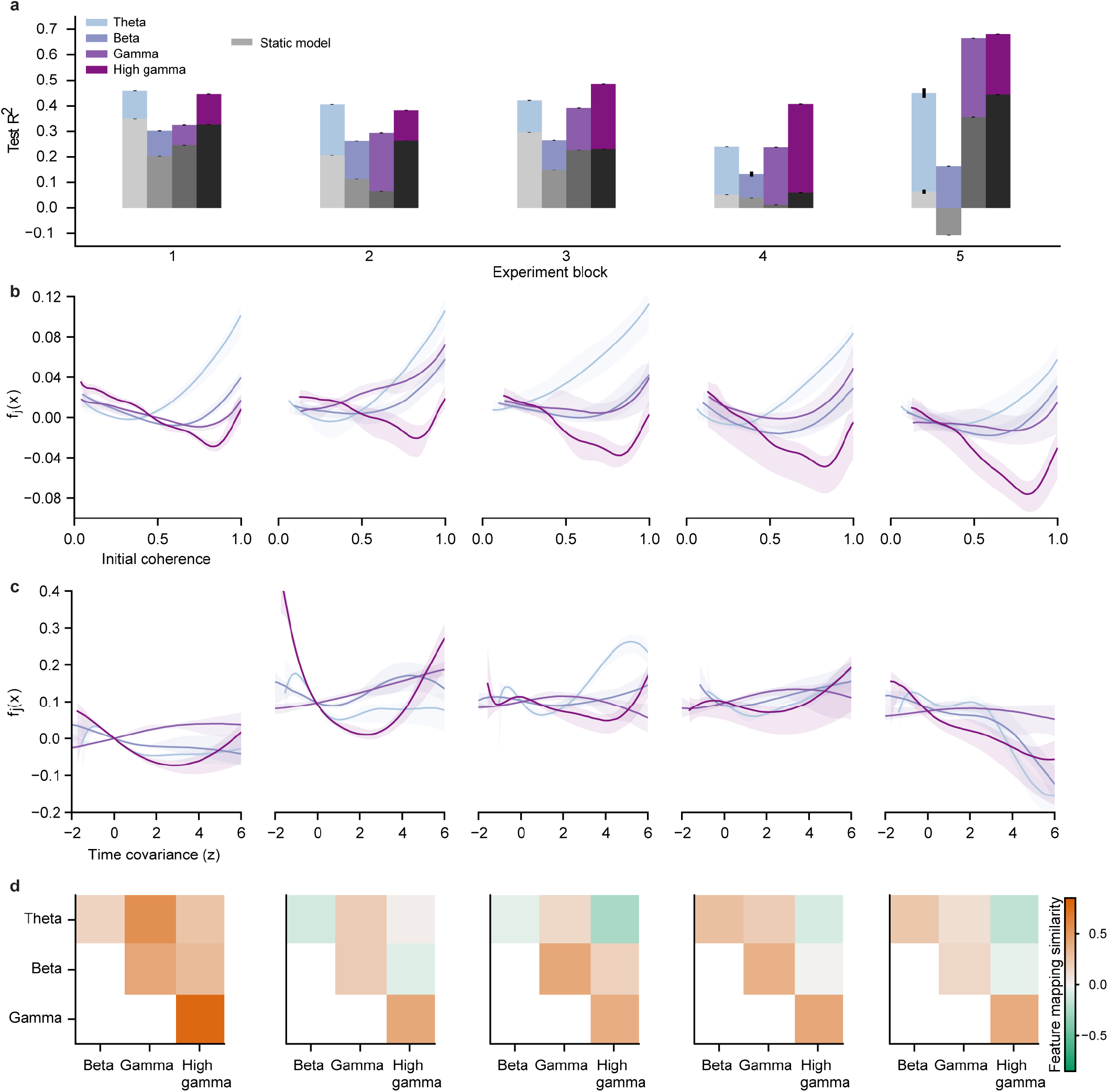
Feature mappings mediating FCC over a network are dependent on the stimulation history. Results in this figure correspond to prediction of SS-FCC; results are similar for prediction of RS-FCC. **a**, Bar graphs indicating r-squared accuracy on held-out data of the time-varying and static model predictions. Greyscale bars indicate the static model while colored bars indicate the time-varying model. The time-varying models outperformed the static model over all frequencies and experiment blocks. Error bars indicate 95% confidence intervals. **b-c**, Example feature mappings of two network features, **b** initial coherence and **c** time covariance (z-scored) used in the time-varying model. We observed that the feature mappings change over successive experiment blocks. Shading indicates 95% confidence intervals. **d**, Heatmaps of similarities of feature mappings between models trained for different frequency bands. Similarity was quantified by feature-averaged cosine similarity of the feature mappings. We observed that while feature mappings were similar at the onset of the stimulation, they diverged over the time-course of stimulation.

## Discussion

Here we leverage advances in neural interface technology and interpretable machine learning to induce and model large-scale FCC. We augment the standard framework of stimulation-induced FCC by considering the effects of the pre-existing network structure and by analyzing FCC over a large-scale (~ 1cm^2^) network. We show that stimulation induces widespread FCC in both the stimulated-state and the resting-state, and that the underlying network structure mediates these effects. Our modeling framework represents a readily translatable method of characterizing the underlying network structure to predict the outcome of stimulation. Our findings explain inconsistent results historically reported *in vivo* (*10*–*12*, *14*–*18*) and pave the way for increased understanding and development of neural stimulation interventions.

### The influence of the stimulation protocol

While brain stimulation can be delivered to a local area, the highly-interconnected nature of the brain ensures that any local region tightly modulates and is modulated by other regions. An analysis of the effects of stimulation which only considers the stimulation protocol thus prioritizes the direct, local effect of stimulation over the mediation of the response by the brain network.

For neural stimulation interventions targeting longer-term FCC *in vivo*, the protocol-focused approach has been found lacking. The dominant framework for stimulation-induced targeted connectivity change was developed in *vitro*, a setting in which the cortical network is largely nonexistent and the stimulation protocol can be extremely precise. Attempts to translate STDP from its native context of monosynaptic connectivity between isolated neuron pairs *in vitro* to functional connectivity between larger-scale neural structures *in vivo* have been largely inconsistent, with some reports of promising results (13) but also of off-target (unstimulated) connectivity changes (*10*, *11*, *14*–*16*) and a lack of response to the protocols (*10*, *11*).

Here, we study real-time network-level FCC as SS-FCC and longer-term FCC as RS-FCC. We found that, in accordance with existing protocol-focused approaches, the stimulation protocol influences both SS-FCC and RS-FCC. However, further analysis indicated that this influence is minor, and that a protocol-focused analysis of stimulation only explains a small fraction of the effects. This is a novel finding, and represents a substantial departure from the standard analysis of the effects of stimulation which prioritizes the stimulation protocol.

Our observation that the stimulation delay was able to predict some RS-FCC between stimulation sites, and that the stimulation delay was one of the top two protocol features for network-wide FCC prediction, is aligned with the delay-focus of STDP-informed connectivity modification approaches (*7*–*10*, *13*–*16*). However, the small amount of variance explained by delay indicates that delay alone is not sufficient for accurate FCC control.

### Network-level analysis of neural stimulation

Here, we model the cortical network as both the mediator and target of the effects of stimulation. Our focus on the network structure is facilitated by our use of novel neurotechnologies for stimulation and recording. We used a large-scale μECoG array to record from a large-scale cortical network. This differed from past studies in which the inclusion of a small set of unstimulated neural sites had primarily been relegated to that of a statistical control (*10*, *14*, *15*). In addition, we used convection-enhanced delivery of the optogenetic virus to obtain widespread opsin expression (*30*, *45*), which allowed us to stimulate many distinct sites of the sensorimotor cortex and to record neural activity during stimulation without artifact.

We estimated a complete network between all recording sites by calculating their pairwise coherences, which is highly correlated to other functional connectivity metrics (*12*, *18*). In addition, unlike metrics such as evoked response, coherence can be calculated in the absence of stimulation. This had the dual advantage of allowing us to estimate connectivity during resting-state blocks and between unstimulated sites. We note that the latter point is especially relevant, as most “network-level” analyses of stimulation only observe a partial network as they do not analyze connectivity between unstimulated regions (*17*, *18*, *22*, *46*).

We then incorporated details of the distributed structure of these networks for prediction of FCC. Recent studies have indicated that pairwise coherence between two sites is correlated to their change in coherence following stimulation (*11*, *17*, *26*). However, any pairwise analysis treats connections as independent from another and does not capture the distributed structure of the network. In order to incorporate details of the distributed network structure into our predictive model, we turned to advances in graph theory and theoretical neuroscience. Graph theory is a branch of mathematics concerned with the study of graphs, and has developed metrics for summarizing and comparing graphs, such as clustering coefficients, measurements of centrality, and graph distances (*36*–*39*). Theoretical neuroscience has similar measures, termed “motifs,” which are used to analyze simulated brain networks (*40*–*43*). Just as simple pairwise connectivity between electrodes describes their first-order connectivity, these measures summarize higher-order, distributed connectivity within a network (*47*). The network features we developed and used in our model were inspired by these existing metrics and similarly encode higher order, distributed connectivity information of the network. For example “coherence with network” and “length 2 path strength” quantify general whole-network connectivity strength between two sites, while “electrode covariance” and “coherence difference” quantify the similarity of connectivity structure to the entire network between two site. Indeed, our model confirmed their relevance by identifying “time covariance” and “coherence difference,” two higher-order connectivity metrics, as the most important features for FCC prediction.

One alternative framework for inducing targeted network-wide FCC is network control theory (*48*), which attempts to use control-theoretic methods to control network activity in real-time. However, it has thus far has been limited to quantifying theoretical controllability of networks (*26*, *46*, *49*) rather than inducing targeted FCC. Here, we validate our framework on the maximally clinically-translatable NHP model and demonstrate that it can accurate predict network-level stimulation-induced FCC.

### The cortical network as the mediator and target of stimulated-state and resting-state functional connectivity changes

Both real-time and long-term modulation of brain networks are relevant for neural stimulation-based medical interventions. Here, we induced, estimated, and modeled network-level FCC which we observed during stimulation as well as after stimulation. The high correlation in FCC and high similarity in feature mappings between the two contexts hint at shared processes underlying SS-FCC and RS-FCC. Practically, this also means that observing stimulation-induced FCC in real-time may lead to more accurate longer-term FCC.

Interventions targeting real-time neuromodulation often do not consider effects beyond local activity modulation. However, this neuromodulation has a secondary effect of altering real-time network synchronization, observed here as network-level SS-FCC, and which persist after stimulation as RS-FCC. As such, the complete effects of these interventions can only be fully understood by augmenting their scope of analysis to include network-level FCC.

Our observation that the underlying network mediates RS-FCC demonstrates that interventions directly targeting RS-FCC must consider the underlying network for accurate induction of connectivity changes. The historical neglect of the underlying network structure in mediating RS-FCC could provide an explanation for reports of off-target connectivity changes and inconsistent connectivity changes between stimulation sites (*10*–*12*, *14*–*16*). We note that, while many of these studies use STDP-informed stimulation which is designed to elicit changes in anatomical connectivity, we cannot pinpoint the nature of our observed FCC as either solely anatomical or activity-based, as functional connectivity is related to both (*50*, *51*).

Finally, our results offer a potential explanation for how underlying network-level functional connectivity shapes the therapeutic response to stimulation. Studies have shown that aberrant functional connectivity underlies many neural disorders (*1*–*3*, *52*), and that neural stimulation alters network-level functional connectivity (*12*, *17*, *18*) and can treat neural disorders (*6*, *31*). As such, therapeutic response to stimulation may be a direct result of a stimulation-induced change from aberrant to healthy network-level functional connectivity. This change may in turn be determined by the underlying network-level functional connectivity, as recent work has indicated that underlying functional connectivity shapes the therapeutic response to stimulation (*23*–*25*), and here we have also shown that it shapes stimulation-induced network-level FCC. Further work which holistically studies the underlying network, neural stimulation, FCC, and neural disorders is needed.

### The evolving response to successive neural stimulation

Our observation that the time-ordered block number was the most important protocol feature indicates that the temporal location of repetitive stimulation is important for consideration. This observation is in accordance with earlier work from our lab (*11*) in which we reported sharp differences in functional connectivity between an initial stimulation block following stimulation blocks. We further investigated this phenomenon with our time-varying model.

With the time-varying model we observed that feature mappings describing the response to stimulation evolve in response to repeated stimulation. This result indicates that stimulation interventions likely need to be adapted over time, as repeatedly stimulating will not repeatedly induce the same response nor even be governed by the same network rules mediating the response. This is consistent with the increased success of some closed-loop neural stimulation interventions over open-loop alternatives (*6*), as it acknowledges that the effects of stimulation can be made more accurate when the stimulation is informed by recent neural activity. However, we additionally show that stimulation should be adapted to characteristics of the whole network, not only to local activity.

### Interpretable predictive modeling beyond the linear paradigm

In this study we capitalized on our network-level treatment of FCC by using a data-driven nonlinear modeling approach. In doing so, we were able to develop an accurate predictive model and estimate feature mappings displaying the relationships between the protocol and network features and FCC. These feature mappings have the dual purpose of elucidating underlying mechanisms of network-level response to stimulation, and informing future stimulation interventions.

Data-driven statistical models often fall into one of two approaches: simplistic linear models or complex “black-box” models. While the simple relationships identified by linear models ensure that the mappings are interpretable, linearity is an assertion, not a property learned from the data, and therefore linear models may not capture the complexity of the true input-output relationships. At the other extreme, prediction-focused black-box models can learn complex input-output mappings at the expense of interpretative ability such as observation of the mappings. The unique modeling approach we employ in our study displays advantages of each as it can identify complex mappings between the features and FCC while allowing the features mappings to be observed. As it is a data-driven and generalized additive model, it retains the ability of identifying linear mappings when such a mapping is supported by the data, while also being able to identify more complex nonlinear mappings when appropriate.

We found that eliminating the artificial and restrictive assumptions of the linear model yielded dramatically improved predictive performance. In particular, as the complexity of our input features increased from simple stimulation parameters to the actual structure of the cortical network, their feature mappings exhibited strong nonlinearities and thus would not be able to be fully represented by a strictly linear approach.

Finally, in light of recent unsuccessful attempts to model the effects of neural stimulation with nonlinear models (*46*), our data-driven method for inferring feature mappings of variable complexity represents a timely contribution for principled and accurate modeling of the effects of brain stimulation.

### Future work

Our neural recordings were limited to a μECoG on the primary sensorimotor cortex surface; further work is needed to verify that our findings are consistent across brain regions, at various cortical depths, and at larger and smaller scales. In addition, as all stimulation experiments consisted of paired optogenetic stimulation, our observations should be verified with other stimulation modalities and protocols to verify their generality. Our method for parsing the network structure, along with the hierarchical additive model, can be easily extended to these cases. If our findings are verified across a broad range of brain regions and stimulation modalities, then this would indicate that existing stimulation-based therapies must factor in the underlying network in order to arrive at a complete understanding of their effects.

Here we have constructed a set of network features which we have shown are relevant for FCC prediction. While some interpretable mappings of interest were described in the results section, others which were more complicated, such as the most-important “time covariance” feature, could not be easily interpreted. Our features broadly encapsulate various spatiotemporal connectivity motifs present in the network connectivity, and as such can be further dissected and parsed for purposes of interpretation. For example, the importance of “time covariance” indicates that the temporal relations of pairwise coherence contains information relevant for FCC prediction; this knowledge can be used to construct more detailed studies with the goal of uncovering the underlying processes governing this relation.

Our finding that the network mediates the plastic response to stimulation can guide the development of stimulation treatments, but does not explicitly offer a prescriptive method for specifying an exact stimulation protocol. Instead, our model acts as an encoder which yields a prediction for the effects of stimulation given a specific stimulation protocol and network state. The creation of a corresponding decoder could be used to directly prescribe a stimulation protocol to elicit desired FCC. Such a model could also be deployed in real-time as a network-informed controller for FCC.

### Neuroscientific and clinical utility of our insights

Here we have shown that the cortical network structure mediates the response to stimulation, both for real-time and longer-term contexts. Our findings provide a possible explanation for how underlying functional connectivity shapes the therapeutic response to stimulation, and for previously reported inconsistent results of neural stimulation. Our framework, only made possible by novel advances in neural interfaces, interpretable machine learning, and graph theory, represents an efficient method for parsing network structure for accurate prediction of stimulation-induced FCC. It can be readily applied to other network-informed approaches of neural stimulation, and can be used to interrogate, improve, and develop novel neural stimulation interventions for neural disorders.

## Materials and Methods

Two adult male rhesus macaques (monkey G: 8 years old, 17.5 Kg; monkey J: 7 years old, 16.5 Kg) were used in this study. All procedures were performed under the approval of the University of California, San Francisco Institutional Animal Care and Use Committee and were compliant with the Guide for the Care and Use of Laboratory Animals.

### Large-scale neural interface

Stimulation and recording of the cortex were achieved using our large-scale optogenetic interface consisting of a semi-transparent micro-electrode array, semi-transparent artificial dura, titanium implant, widespread optogenetic expression, and laser-delivered optical stimulation (*30*). Neurons in the primary sensorimotor cortex were optogenetically photosensitized via viral-mediated expression of the C1V1 opsin. We injected a viral cocktail of AAV5-CamKIIa-C1V1(E122T/E162T)-TS-eYFP-WPRE-hGH (2.5 × 10^12^ virus molecules/ml; Penn Vector Core, University of Pennsylvania, PA, USA, Addgene number: 35499) in the primary sensory (S1) and primary motor (M1) cortices of the left hemisphere of the two rhesus macaques using the convection-enhanced delivery technique as described in (*30*, *53*). We infused 200 μL of the cocktail was over four sites in monkey G and 250 μL over five sites in monkey J.

The chronic neural interface was implanted by performing a 25mm craniotomy over the primary sensorimotor cortices of the left hemisphere of two rhesus macaques, replacing the dura mater beneath the craniotomy with a chronic transparent artificial dura, and attaching a titanium cap over the craniotomy. During experiments we removed the artificial dura and placed a custom 92mm^2^ micro-electrocorticography array of 96 electrodes on the cortical surface for recording neural activity. The μECoG arrays consisted of platinum-gold-platinum electrodes and traces encapsulated in Parylene-C (*54*).

Neural data was recorded by sampling local-field potentials at 24 kHz from the μECoG array using a Tucker-Davis Technologies system (FL, USA). We stimulated the cortex by delivering light via a fiber optic (core/cladding diameter: 62.5/125 μm, Fiber Systems, TX, USA) connected to a 488 nm laser (PhoxX 488-60, Omicron-Laserage, Germany) positioned above the array such that the tip of the fiber-optic cable touched the array.

The interface used for the subjects in this study is further described in (*30*). Of the dataset used for this study, a subset overlaps with the dataset used in our previous publication (*12*).

### Verification of optogenetic expression and activation

We verified expression of the C1V1 opsin by fluorescent imaging of the eYFP marker (Fig. S4), and by recording optogenetically-evoked neural responses (Fig. S5), as described in (*30*). We observed widespread fluorescence in the primary sensorimotor cortices (Fig. S4). We also observed neural activity elicited by illumination site, propagating within the stimulated anatomical region and across the central sulcus (Fig. S5). Also, as detailed in the “Signal preprocessing” section, at the beginning of each experimental session we verified that recorded activity was not due to photoelectric artifacts by stimulating at 500 Hz and confirming that we did not see matching LFP traces. We also verified that the evoked neural responses were due to optogenetic activation and not another light-induced factor such heating by illuminating outside the sites expressing fluorescence and recording neural activity. We found that such illumination did not evoke neural activity, confirming that the observed neural activation from our experimental sessions was due to optogenetic activation.

### Structure of experimental sessions

Throughout each experimental session, subjects were watching cartoons while seated in a primate chair and headfixed. At the beginning of an experiment the chamber was opened, the artificial dura was removed, and the μECoG array was placed on the cortical surface. Throughout the experiment, the exposed cortex was irrigated with saline to ensure that it remained moist. At the end of each experiment, the μECoG array was removed from the cortical surface, the artificial dura was replaced on the brain, and the chamber was closed.

We performed a total of 36 experimental sessions over multiple days and two subjects. The experimental sessions consisted of recording while no stimulation took place, which we termed a “resting-state block,” and recording during stimulation, which we termed a “stimulation block.” Each resting-state block was 5 minutes long and each stimulation block was 10 minutes long. Each experimental session consisted of alternating resting-state blocks and stimulation blocks, with 6 total resting-state blocks and 5 total stimulation blocks, as a resting-state block was completed before the first and after the final stimulation block. The μECoG array was replaced on the cortex for each experimental session. Additionally, each recording block of an experimental session had the same stimulation protocol.

We stimulated two locations per experimental session which varied between experimental sessions but remained fixed throughout an experimental session. The locations were randomly chosen across the array. Optogenetic activation at the chosen stimulation site was verified before beginning the experimental session, by stimulating and observing activation in the LFP traces. We ensured that these traces were due to neural activation and not photoelectric artifact by the method described in the “Signal preprocessing” section. We imaged the stimulation locations on the array and subsequently identified the closest recording electrode to each stimulation location. The distance from stimulation site to the nearest electrode was no more than 500 μm.

We applied stimulation using a paired-pulse protocol of 5 ms pulses separated by a session-specific delay (Fig. 1). A pulse width of 5 ms was chosen because it elicited reliable responses in M1 and S1, and has a close duration to previously used targeted plasticity protocols (*13*). The session-specific delays were randomly chosen at the beginning of each experimental session to be 10 ms, 30 ms, or 100 ms; once chosen they remained fixed for an experimental session. These values were motivated by previous studies reporting that 10 ms and 30 ms delays induced reliable connectivity changes while 100 ms delays did not induce changes (*9*, *13*). The paired-pulses were repeated every 200 ms for the entirety of each stimulation block; this period was chosen to allow for enough time for the chosen delays while maximizing the number of paired-pulse stimulations within a conditioning block.

We performed a total of four additional control sessions over 3 days and two subjects. The structure of the sessions remained identical to the experimental sessions described in this section, except that no stimulation took place.

### Signal preprocessing

Before beginning an experimental session, we tested for photoelectric artifact. This was done by stimulating at frequencies faster than C1V1 off-kinetics (eg. 500 Hz) (*29*) and examining the LFP traces. If this high frequency stimulation elicited response in the LFP traces then this was deemed to be photoelectric artifact, and the illumination site was moved to another location (*54*).

Before recording, electrode impedances were measured and those with high impedance were excluded from further analysis. We also examined the broadband recorded surface potentials and excluded those with low signal-to-noise ratio. The total amount of data removed constitutes no more than 15% of the raw time series. We then downsampled the data to 1 kHz after applying a low-pass Chebychev filter for anti-aliasing.

### Signal processing to obtain a time-varying coherence network

Within each stimulation block and each resting-state block, the recorded LFP were partitioned into non-overlapping 20-second windows. All quantities computed from the raw LFP time series were computed on a per-window basis, then summarized across all windows in the block.

We computed an estimate of the coherence between all non-faulty electrodes for each 20-second window in every resting-state block and stimulation block. We denote by *E* the number of non-faulty electrodes and denote the multivariate LFP time series for a given window by the two-dimensional array 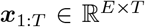, which consists of the time-ordered concatenation of the LFP recordings ***x***_*t*_ = (*x*_1*t*_,…,*x_Et_*)^*T*^ for each electrode at time *t*. We estimated the multivariate spectral density function 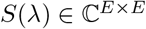 for the time series ***x***_1:*T*_ by Welch’s method (*55*). The coherence between electrodes *i* and *j* at frequency *λ* is denoted as *C_ij_*(*λ*) and computed from the estimated spectral density via *C_ij_*(*λ*) = |*S_ij_*(*λ*)|^2^/*S_ii_*(*λ*)*S_jj_*(*λ*). We computed the coherence at 403 linearly spaced frequencies in the interval (0, 200) Hz.

Summaries over four frequency bands, 4-7 Hz (Theta), 12-30 Hz (Beta), 30-70 Hz (Gamma), and 70-199 Hz (High Gamma), were computed by averaging the coherence estimates within each band. Thus, for each session, the raw LFP time series was transformed into a sequence of matrix-valued quantities representing the average band-limited coherence in each 20-second interval. Since the coherence is symmetric and the diagonal conveys no information on pairwise behavior of the LFP electrode signals, we retained only values in the upper triangle above the diagonal for modeling and prediction.

We defined SS-FCC as the difference between the mean coherence of an electrode pair calculated during a stimulation block and the preceding resting-state block. We defined RS-FCC as the difference between the mean coherence of an electrode pair calculated during a resting-state block and the preceding resting-state block. We use FCC as a general term to refer to either SS-FCC or RS-FCC. Let 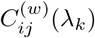 be the coherence between electrodes *i* and *j* at frequency *λ_k_* in the 20-second window *w*, as estimated by the procedure described above. The band-limited coherence was obtained by averaging over the indices *k* ∈ *K_b_* corresponding to frequency band 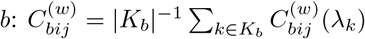. Within an experimental session, we denote by *W_R_ℓ__*, *ℓ* ∈ {1, 2, 3, 4, 5, 6} and *W_S_ℓ__*, *ℓ* ∈ {1, 2, 3, 4, 5}, the collections of 20-second windows belonging to resting-state block *ℓ* or stimulation block *ℓ*, respectively. Note that the additional resting-state block corresponds to a final recording block after the 5th stimulation block. The electrode coherences were summarized for each of these blocks by their mean values:

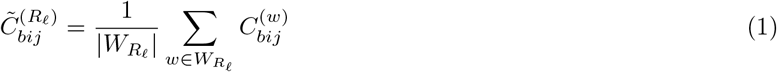

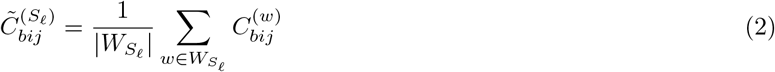

The SS-FCC was then computed as

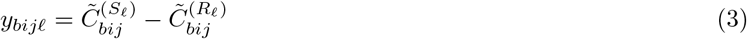

and the RS-FCC was computed as

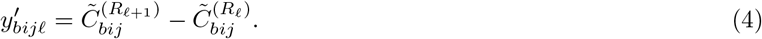

### Nonlinear modeling of stimulation-induced coherence change

We used a nonlinear additive model to explain the observed FCC in terms of features derived from both the experimental protocol and the information on functional connectivity available in the resting-state block prior to stimulation. In this framework, a single observation was denoted by the pair (*y_n_, **x**_n_*), *n* = 1,…, *N*, with 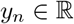 an FCC measurement and 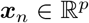 the corresponding features. In switching to the single index *n* we indicate that the data was aggregated over all unique electrode pairs and all experimental blocks. The dependence on the band *b* was subsequently omitted for notational simplicity; the same analysis was repeated for the data in each frequency band. The FCC was modeled as

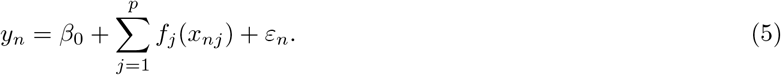

The model consists of an intercept *β_0_*, nonlinear functions *f_j_* controlling the impact of each feature *j* = 1,…,*p* on the response, and an error term *ε_n_*.

The utility of the model derives from the nonlinearity of the feature mappings and the additive procedure through which they are combined to yield a prediction for the FCC. The additive structure of the model allows for the visualization and interpretation of individual feature-response relationships. For each feature *j* = 1,…, *p*, this is represented by the feature mapping *f_j_*. Moreover, the predictive impact of feature *j* can be measured by the decrease in prediction error on a test set relative to a model in which the feature is omitted (or equivalently, a full model is estimated while enforcing *f_j_* = 0). The same analysis can be applied to groups of features. While additivity is essential for model investigation, the nonlinearity of the feature mappings enables the identification of more complex relationships than can be expressed in a linear model. Linearity is a strong modeling assumption that is both difficult to justify scientifically and, as demonstrated in the results, harmful in terms of predictive accuracy.

### Feature representation of processed data

A single observation of the FC corresponds to a unique pair of electrodes and a specific recording block. The corresponding features were constructed to satisfy two objectives: first, that the subsequent analysis separates the influence of the protocol parameters from that of the network structure of the functional connectivity; second, that all information in the features was available prior to stimulation.

To satisfy the first objective, we partitioned the features into two groups: *protocol* features, which summarize aspects of the experimental setting and protocol; and *network* features, which summarize information in the electrode coherences during baseline recording. The protocol features describe the key parameters of the experimental framework. The designation “network features” derives from consideration of the estimated band-limited coherence as the adjacency matrix of an undirected, edge-weighted graph. The network features were intended to serve as simple but informative summaries of the connectivity information available in recordings of baseline activity prior to stimulation. They correspond to summary statistics computed over spatial or temporal ranges, basic quantities pertaining to the estimated spectrum (*56*), or network features that have found previous application in graph analysis (*36*, *57*). Finally, a subject-level indicator was included to adjust for potential global differences in FCC between the two macaque subjects. The adjustment is included in all models, as the subject is neither an aspect of the stimulation protocol nor a property of the baseline connectivity network.

#### Protocol features

The protocol features were comprised of the categorical measurements `Anatomical region, Delay`, and `Block number`, as well as the real-valued features `Distance, Stim1 distance to closer, Stim1 distance to further, Stim2 distance to closer`, and `Stim2 distance to further. Anatomical region` indicates whether the two electrodes corresponding to a given measurement are both in region M1, both in region S1, or if one is in each region. `Delay` encodes three levels of time-delay in the pulse of the paired laser stimulation: 10, 30, or 100 ms. While the delay parameter in the stimulation protocol is real-valued, in the context of our data it is only measured at three distinct settings. We therefore chose to estimate an effect for each setting individually rather than to estimate a nonlinear function of the delay given observations at only three points. `Block number` indicates the time-order position of the experimental block in which the observation was recorded. In the “block analysis” regression design, we removed this feature and instead allowed all feature mappings to vary with the block number, thus investigating the predictive impact of allowing the entire model to evolve dynamically over the discrete time-stages of an experimental session. `Distance` measures the distance between the two electrodes. `Stim1 distance to closer, Stim1 distance to further, Stim2 distance to closer`, and `Stim2 distance to further`, encode distances between the stimulation sites and the closer and further electrodes of the electrode pair being predicted.

#### Network features

The network features were computed from the LFP time series recorded during the resting-state block preceding the stimulation for which FCC are being predicted. Except for the `Phase` feature, all of these features were derived from the tensor obtained by time-concatenation of the coherence matrices across all windows in the resting-state block.

We introduce some tensor notation prior to defining these features. The coherence tensor for a given resting-state period is a three-dimensional array, denoted as 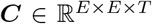. Two-dimensional slices through the array at time *t* or electrode *i* are indicated as *C_t._* and *C*·*i*·, respectively. One-dimensional vectors obtained by fixing an electrode *i* and a time point *t* are denoted 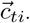; vectors obtained by fixing both electrodes *i* and *j* are denoted 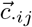. The scalar coherence value at time t for electrodes *i* and *j* is denoted *c_tij_*.

The `Initial coherence` indicates the mean coherence between electrodes *i* and *j* at baseline:

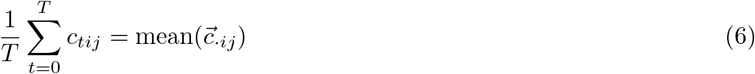

The `Coherence with network` summarizes the average coherence between electrodes *i* and *j* and the remaining electrodes in the array. This corresponds to the normalized sum of vertex strengths of nodes *i* and *j* (*36*):

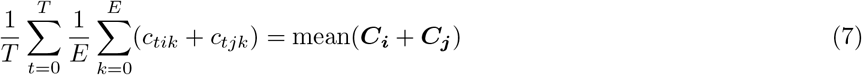

The `Coherence difference` measures the mean absolute difference in coherence between electrodes *i* and *j* to other electrodes in the network:

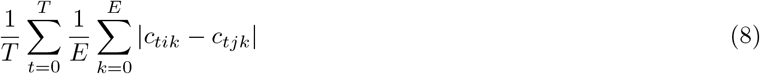

The `Length 2 path strength` represents the average strength of length-2 paths connecting electrodes *i* and *j*. This feature is similar to standard graph clustering coefficient metrics quantifying the total edge-weight of triangle motifs that include a given node (*36*, *37*). However, it excludes the direct edge between electrodes *i* and *j*, which reduces correlation with the initial coherence. Alternatively, the `L=2 path strength` can be considered the unnormalized cosine similarity between the network-coherence vectors of electrodes *i* and *j* (*57*):

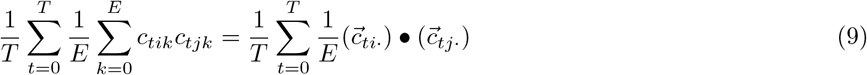

The `Coherence with stim sites` is the average coherence between electrodes *i* and *j* to the two optogenetic stimulation sites. Below, the indices *a* and *b* refer to the electrode sites corresponding to the laser locations for a given experiment:

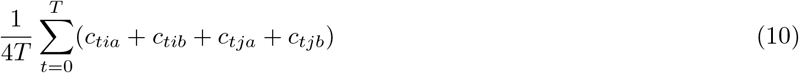

The `Electrode covariance` and `Time covariance` capture the variability of the coherence measurements over the electrode array and over the total number of 20-second time windows in a resting-state block, respectively.

The `Electrode covariance` represents a time-average of the covariance between the vectors representing the connectivity of electrodes *i* and *j* to the rest of the network:

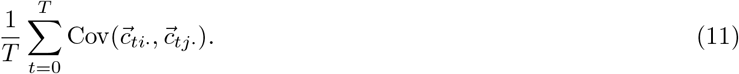

The `Time covariance` is the average across all other electrodes *k* of the covariance between the time series of coherence values to electrodes *i* and *j*:

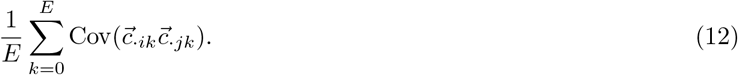

Finally, the `Phase` was the only network feature not derived from the coherence tensor, since the coherence only contains magnitude information from the estimated spectral density. Writing the (*i*, *j*)^*th*^ cross-spectral component computed within window *w* as

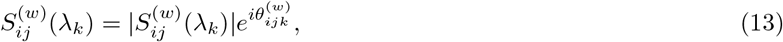

and denoting by *K_b_* the set of indices *k* such that *λ_k_* is in frequency band *b*, the phase feature *θ_ij_* in block *ℓ* is given by

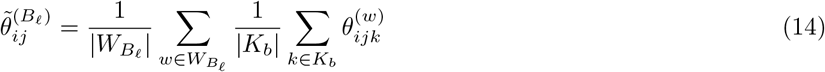

##### Outlier removal

Nonlinear regression methods are sensitive to extreme outliers, which can exert highly disproportionate influence on the model estimate. We wished to achieve robustness against such outliers while minimizing perturbation of the data and subsequent analysis. Therefore we adopted a highly permissive definition, by which an observation was considered an outlier if its absolute deviation from the mean exceeded 20 times the interquartile range along any dimension (i.e. feature). Across all frequency bands and models, the maximum amount of data excluded by this procedure was less than 0.1% of the total number of observations.

#### Regression designs

We investigated two different configurations of the regression design matrix within our nonlinear modeling framework. For each design, the data were aggregated across all of the experimental sessions for each subject. While significant session heterogeneity suggested that even stronger prediction results were possible for models fit to individual sessions, our objective was to estimate a model that generalized well in the sense of accurate prediction on data from many sessions.

##### Full data

First we investigated the performance of the nonlinear model on the full data, which included all experimental blocks from all sessions of both subjects. This yielded *N* = 481, 505 observations of *p* = 16 features, which expanded to dimension 21 after dummy-coding of the categorical variables `Delay, Region, and Block number`, and to dimension 22 after inclusion of the categorical animal subject indicator.

##### Block interactions

Under the full data design, the time evolution of the response was limited to global shifts corresponding to the categorical feature `Block number`. We subsequently investigated the impact of allowing the shape of the feature mappings to vary with the block number, thus expanding the model’s capacity to capture time variation in feature-response relationships over the course of repeated stimulation. This constituted an interaction design, whereby the categorical feature `Block number` was removed, and the remaining features were each augmented with interaction terms for binary indicators that denoted whether the measurement was made in each of blocks 2, 3, 4, and 5. The number of observations remained *N* = 481, 505, while the number of features expanded to *p* = 94.

#### Estimation of the nonlinear additive model

We used a train-test paradigm to assess the predictive performance of the nonlinear model on unseen data. For each frequency band and regression design, the data were split uniformly at random such that 70% of the observations were assigned to the training set and the remaining 30% were assigned to the test set. All model parameters and hyperparameters were selected using data from the training set only. Prior to estimation of the model parameters, the real-valued features were standardized such that they have mean zero and unit standard deviation. Whenever the data were split between train and test sets, the parameters of the linear transformation corresponding to standardization were computed from the training data and applied to the test data.

The nonlinear feature mappings *f_j_*, *j* = 1,…, *p*, in Eq. (5) were represented in the basis of polynomial functions

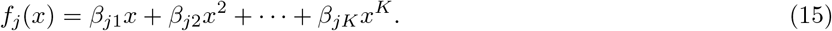

The complete set of parameters to be estimated was thus ***β*** = (*β*_0_, *β*_1_,…, *β_p_*), where 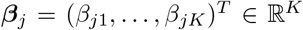. The *order* of the polynomial representation for *f_j_* is defined as the maximum integer value *k* ∈ 1,…, *K* for which *β_jk_* is nonzero. The smoothness of *f_j_* is determined by the order *k* of its polynomial representation and the magnitude of the coefficients *β*_*j*1_,…, *β*_*jk*_. Numerical implementation requires the specification of an upper bound *K*, corresponding to truncation of the infinite basis at some maximum order. We used *K* = 10 in all experiments; this decision was justified by the fact that the parameter estimates decay either to negligible values or to exactly zero before reaching order *K* ≤ 10, as a result of the hierarchical penalization of the model coefficients.

All categorical features were dummy-coded such that the *C* categories for each feature are represented by *C* – 1 indicator variables. Polynomial expansion of these variables generates identical columns and thus rank-deficiency in the design matrix, which leads to instability in the optimization routine. This problem was avoided by truncating the polynomial expansion of all categorical variables to order 1 rather than order *K*. There is no resulting loss of generality from this truncation. Rather, this expanded the framework to allow for simultaneous estimation of smooth nonlinear functions of continuous features and discrete functions (that can be visualized for example, as a step function) of categorical features.

#### Automatic order selection via hierarchically penalized estimation

Given an observation of the features ***x***, the model prediction of the SS-FCC or RS-FCC is given by

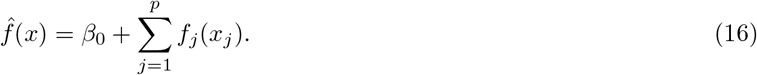

We used the standard square loss on the training set *D^train^*

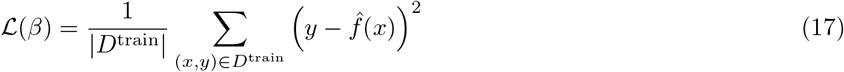

as the model goodness-of-fit criterion. Direct optimization of model fit on a training set over a sufficiently complex model class is well-known to yield estimates that perform poorly on unseen data, an issue commonly known as overfitting. We avoided this problem by adoption of a penalized estimation framework specifically designed to control the magnitude of the parameter estimates and automatically select the order of each feature mapping in the model. This approach augments the standard square loss with a penalty term Ω_*j*_(*β_j_*; *α*, *λ*) for each feature *j*, defined as

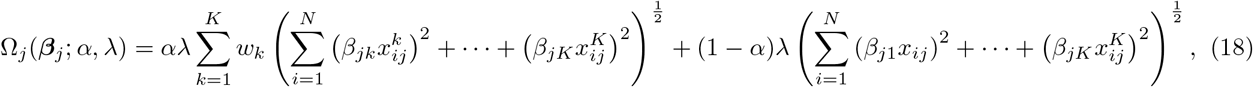

with hyperparameters 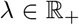 and *α* ∈ [0, 1]. The the weights *w_k_* = *k*^3^ – (*k* – 1)^3^ were chosen to satisfy theoretical criteria that guarantee statistical convergence of the estimator over a class of smooth functions (*28*).

The model parameters were estimated by solving

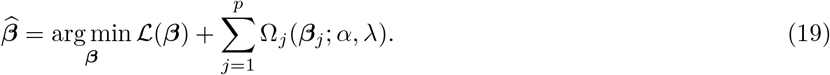

We used a block coordinate descent algorithm as implemented in the R package `HierBasis` (*28*).

The first term in the penalty Ω_*j*_(*β_j_*; *λ*, *α*) induces hierarchical sparsity in the estimate of *β_j_* = (*β*_*j*1_,…, *β*_*jK*_)^*T*^, while the second term in (18) shrinks the magnitudes of the estimated coefficients towards zero. Hierarchical sparsity guarantees that if an estimated coefficient 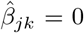, then all higher-order coefficients for that feature 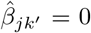, with *k* < *k*’ ≤ *K*; this is equivalent to selecting an order *k* – 1 representation for the feature mapping *f_j_*. The selection procedure is thus *automatic* and *data-driven* in that it emerges as a direct mathematical consequence of the penalized estimation framework, which seeks the best model fit to the data under the structural constraints imposed by Ω_*j*_(*β*).

#### Cross-validation for model hyperparameters

The penalized loss function requires specification of two regularization hyperparameters 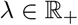 and *α* ∈ [0, 1], which control the overall regularization strength and tradeoff between terms inducing hierarchical and feature-wise sparsity, respectively. The parameter *α* was selected over the range from 0 to 1, inclusive, in increments of 0.1. The parameter *λ* was selected over 100 log-linearly-spaced values between λ_max_ and λ_max_ × 10^-5^.

The value λ_max_ was selected such that all estimated feature mappings are equal to zero for every value of investigated *α*; it was obtained by a backtracking line search algorithm (*58*). This algorithm requires a gross upper bound λ_0_, defined as any value of λ > 0 such that 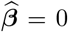 for all grid values of *α*. It was found manually; we found that λ_0_ = 0.1 sufficed for all designs and frequency bands.

We selected the pair (α*, λ*) by first finding the (α, λ) pair minimizing the average *R*^2^ on the validation set over 5-fold cross-validation on the training data. We selected α* as the α-coordinate of this pair, and selected λ* as the largest value of λ such that the mean validation *R*^2^ of (α*, λ) was within one standard error of the mean validation *R*^2^ at (α*,λ*). This follows the “one standard error” strategy for one-dimensional cross-validation (*35*) and represents a conservative approach to regularization corresponding to our preference for smoother feature mapping estimates.

##### Performance measure

We quantified model performance by predictive accuracy on the test set, as measured by the coefficient of determination,

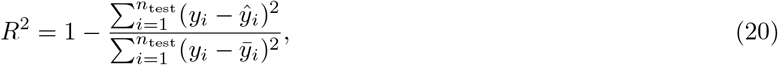

where *ŷ_i_* was the model prediction after cross-validation and estimation on the training set and *ȳ_i_* was the mean of the regression target on the test set.

### Stability, importance, and similarity of learned feature mappings

#### Resampling for feature stability

We took a resampling approach to assess the stability of the estimated feature mappings of the nonlinear model. Following the subsampling heuristic of (*44*), we drew a subsample of size ⌊*N*/2⌋ from the data without replacement and ran the full estimation procedure, including hyperparameter selection, on the subsampled data. The procedure was repeated for 100 independent trials, resulting in 100 estimated functions for each feature. The feature-wise variability in the nonlinear additive feature mappings was then assessed pointwise, by computing upper and lower quantiles for the range of function values at each point in a fine grid.

#### A predictive measure of feature importance

Traditional tests of statistical significance are not available for the coefficients of the nonlinear additive model as the asymptotic distribution of the penalized estimator is not known. Instead, we assessed importance of a feature or group of features by evaluating the difference in predictive performance on the test set between the full model and a model fit without the parameter or parameter group of interest. Removing the feature or feature group is equivalent to estimating the full model under the constraint that their corresponding parameters are all equal to zero. The comparison can thus be viewed as between the full model and one in which a hypothesis of null response is enforced for the feature or group of interest.

Quantifying feature importance via the step-down procedure accounts for the fact that the explanatory power of the feature of interest may partially overlap with other included features. We are thus asking, “what is the marginal explanatory power of this feature, beyond what could be explained by the remaining features in the model?” In the context of our work, this is used to understand, for example, whether the network features provide any additional information over the protocol features. A step-up procedure could not answer such a question.

When estimating the reduced model, we used the regularization parameters (α*,λ*) obtained by cross-validation on the full data.

#### Quantifying feature similarity across frequency bands

Let 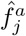 and 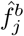 be the estimated feature mappings for feature *j* on the data corresponding to frequency bands *a* and *b*, respectively. We computed a quantitative measure of their similarity,

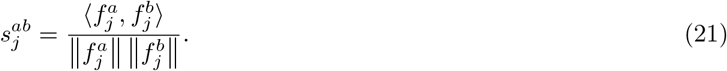

The quantity 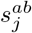 is the cosine similarity of the estimated feature mappings, considered as elements of a common Hilbert space of functions. By definition, 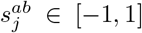. Similarity increases as 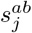 approaches 1 or −1 (which indicates 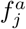 and 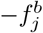 are perfectly similar) and is minimized at 0.

The inner product 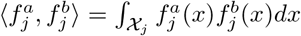 and norm ||*f_j_*|| = ⟨*f_j_*, *f_j_*⟩ are given by integrals whose domain 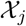 depends on the feature *j*. For real-valued features, we took 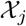 to be the interval [−5, 5], which after standardization corresponds to the range of all observed measurements within 5 standard deviations of the mean. Due to the polynomial representation of the nonlinear feature mappings, these could be computed exactly.

## Acknowledgements

We thank Daniel Silversmith for his help with data collection and Philip Sabes for his laboratory in which the data was collected. This work was supported by the Big Data for Genomics and Neuroscience Training Grant (NIH 5T32LM012419, JB and AGT), the Center for Neurotechnology (NSF ERC 1028725, JB and ESB), the Washington National Primate Research Center (NIH P51 OD010425, AYS), the Eunice Kennedy Shriver National Institute of Child Health and Human Development (NIH K12HD073945, AYS), NSF grant DMS-2023239 (ZH), NSF grant DMS-1722246 (AS), NSF grant DMS-1915855 (AS), and NIH grant R01GM114029 (AS).

## Author Contributions

JB and AGT drafted the manuscript. JB and AYS conceptualized the study. AYS performed the experiments. JB processed the data. AGT performed the nonparametric modeling. All authors revised and edited the manuscript.

## Data and Code Availability

The data supporting the findings of this study can be accessed at the following url: doi.org/10.6084/m9.figshare.16625726. The code supporting the findings of this study can be accessed at the following url: bitbucket.org/yazdanlab/network_level_fcc_modeling/src/master/.

## References

1. A. G. Garrity et al., American Journal of Psychiatry, ISSN: 0002953X (2007).

2. Y. Nakai et al., Scientific Reports 11, 1–10, (https://doi.org/10.1038/s41598-021-81207-6) (2021).

3. C. J. Stam, B. F. Jones, G. Nolte, M. Breakspear, P. Scheltens, Cerebral Cortex 17, 92–99, ISSN: 10473211 (2007).

4. M. D. Fox, M. A. Halko, M. C. Eldaief, A. Pascual-Leone, NeuroImage 62, 2232–2243, ISSN: 10538119, (http://dx.doi.org/10.1016/j.neuroimage.2012.03.035) (2012).

5. B. Sehm et al., Journal of Neurophysiology 108, 3253–3263, ISSN: 00223077 (2012).

6. M. A. Edwardson, T. H. Lucas, J. R. Carey, E. E. Fetz, Experimental Brain Research 224, 335–358 (2013).

7. G. Bliss, T.V.P. and Collingridge, Nature 361, 31–39 (1993).

8. H. Markram, J. Lübke, M. Frotscher, B. Sakmann, Science 275, 213–215, ISSN: 00368075 (1997).

9. G.-q. Bi, M.-m. Poo, J. Neurosci 18, 1–9, ISSN: 0270-6474 (1998).

10. S. C. Seeman, B. J. Mogen, E. E. Fetz, S. I. Perlmutter, The Journal of Neuroscience 37, 1935–1949, ISSN: 0270-6474 (2017).

11. J. A. Bloch et al., Proceedings of the Annual International Conference of the IEEE Engineering in Medicine and Biology Society, EMBS, 6446–6449, ISSN: 1557170X (2019).

12. A. Yazdan-Shahmorad, D. B. Silversmith, V. Kharazia, P. N. Sabes, Elife 7, e31034 (2018).

13. A. Jackson, J. Mavoori, E. E. Fetz, Nature 444, 56–60, ISSN: 14764687 (2006).

14. J. M. Rebesco, I. H. Stevenson, K. P. Körding, S. A. Solla, L. E. Miller, Frontiers in Systems Neuroscience 4, 1–15, ISSN: 16625137 (2010).

15. J. M. Rebesco, L. E. Miller, Journal of Neural Engineering 8, ISSN: 17412560 (2011).

16. W. Song, C. C. Kerr, W. W. Lytton, J. T. Francis, PLoS ONE 8, ISSN: 19326203 (2013).

17. C. J. Keller et al., Journal of Neuroscience 38, 5384–5398, ISSN: 15292401 (2018).

18. Y. Huang et al., The Journal of neuroscience 39, 6122–6135, ISSN: 15292401 (2019).

19. D. Momi et al., NeuroImage 229, 117698, ISSN: 10959572, (https://doi.org/10.1016/j.neuroimage.2020.117698) (2021).

20. E. A. Solomon et al., Nature Communications 9, 1–13, ISSN: 20411723, (http://dx.doi.org/10.1038/s41467-018-06876-w) (2018).

21. K. C. Fox et al., Nature Human Behaviour 4, 1039–1052, ISSN: 23973374, (http://dx.doi.org/10.1038/s41562-020-0910-1) (2020).

22. C. J. Keller et al., Proceedings of the National Academy of Sciences of the United States of America 108, 17234, ISSN: 10916490 (2011).

23. A. Horn et al., Annals of Neurology 82, 67–78, ISSN: 15318249 (2017).

24. J. R. Younce et al., Movement Disorders 36, 662–671, ISSN: 15318257 (2021).

25. M. D. Fox, R. L. Buckner, M. P. White, M. D. Greicius, A. Pascual-Leone, Biological Psychiatry 72, 595–603, ISSN: 00063223, (http://dx.doi.org/10.1016/j.biopsych.2012.04.028) (2012).

26. A. N. Khambhati et al., Network Neuroscience 3, 848–877, (http://dx.doi.org/10.1162/netn_a_00089) (2019).

27. E. S. Boyden, F. Zhang, E. Bamberg, G. Nagel, K. Deisseroth, Nature Neuroscience 8, 1263–1268, ISSN: 10976256 (2005).

28. A. Haris, A. Shojaie, N. Simon, Biometrika 106, 87–107 (2019).

29. R. Prakash et al., Nature Methods 9, 1171–1179, ISSN: 15487091 (2012).

30. A. Yazdan-Shahmorad et al., Neuron 89, 927–939, ISSN: 10974199 (2016).

31. A. M. Lozano, N. Lipsman, Neuron 77, 406–424, ISSN: 08966273 (2013).

32. R. M. Levy et al., Neurorehabilitation and Neural Repair 30, 107–119, ISSN: 15526844 (2016).

33. L. I. Zhang, H. W. Tao, C. E. Holt, W. A. Harris, M. M. Poo, Nature, ISSN: 00280836 (1998).

34. S. Song, K. D. Miller, L. F. Abbott, Nature Neuroscience, ISSN: 10976256 (2000).

35. J. Friedman, T. Hastie, R. Tibshirani, The Elements of Statistical Learning (Springer Series in Statistics New York, 2001), vol. 1.

36. A. Barrat, M. Barthélemy, R. Pastor-Satorras, A. Vespignani, Proceedings of the National Academy of Sciences of the United States of America 101, 3747–3752 (2004).

37. J. P. Onnela, J. Saramäki, J. Kertész, K. Kaski, Physical Review E - Statistical, Nonlinear, and Soft Matter Physics 71 (2005).

38. R. Milo et al., Science, ISSN: 00368075 (2002).

39. R. Milo et al., Science, ISSN: 00368075 (2004).

40. V. Pernice, B. Staude, S. Cardanobile, S. Rotter, PLoS Computational Biology 7, ISSN: 1553734X (2011).

41. L. Zhao, B. Beverlin, T. Netoff, D. Q. Nykamp, Frontiers in Computational Neuroscience 5, 1–16, ISSN: 16625188 (2011).

42. Y. Hu, J. Trousdale, K. Josić, E. Shea-Brown, Physical Review E - Statistical, Nonlinear, and Soft Matter Physics, ISSN: 15502376 (2014).

43. G. K. Ocker et al., Current Opinion in Neurobiology, ISSN: 18736882 (2017).

44. N. Meinshausen, P. Bühlmann, Journal of the Royal Statistical Society: Series B (Statistical Methodology) 72, 417–473 (2010).

45. K. Khateeb, D. Griggs, P. N. Sabes, A. Yazdan-Shahmorad, Journal of visualized experiments: JoVE 147 (2019).

46. Y. Yang et al., Nature Biomedical Engineering (2021).

47. A. R. Benson, D. F. Gleich, J. Leskovec, Science 353, 163–166, ISSN: 00107514, arXiv: 0405123 (2016).

48. D. S. Bassett, O. Sporns, Nature Neuroscience 20, 353–364, ISSN: 15461726 (2017).

49. S. F. Muldoon et al., PLoS computational biology 12, e1005076, ISSN: 1553-7358 (2016).

50. Z. Wang et al., Neuron 78, 1116–1126, ISSN: 08966273 (2013).

51. A. B. Waites, A. Stanislavsky, D. F. Abbott, G. D. Jackson, Human Brain Mapping 24, 59–68, ISSN: 10659471 (2005).

52. R. H. Kaiser, J. R. Andrews-Hanna, T. D. Wager, D. A. Pizzagalli, JAMA Psychiatry 72, 603–611, ISSN: 2168622X (2015).

53. A. Yazdan-Shahmorad et al., Journal of Neuroscience Methods 293, 347–358 (2018).

54. P. Ledochowitsch et al., Journal of Neuroscience Methods 256, 220–231 (2015).

55. P. Welch, IEEE Transactions on audio and electroacoustics 15, 70–73 (1967).

56. P. J. Brockwell, R. A. Davis, S. E. Fienberg, Time Series: Theory and Methods (Springer Science & Business Media, 1991).

57. G. Salton, M. McGill, Introduction to Modern Information Retrieval (McGraw-Hill, Inc., Auckland, 1983).

58. S. Boyd, L. Vandenberghe, Convex Optimization (Cambridge University Press, 2004).

