## Supplement for "Cortical network structure mediates response to stimulation: an optogenetic study in non-human primates"

### Supplementary Materials

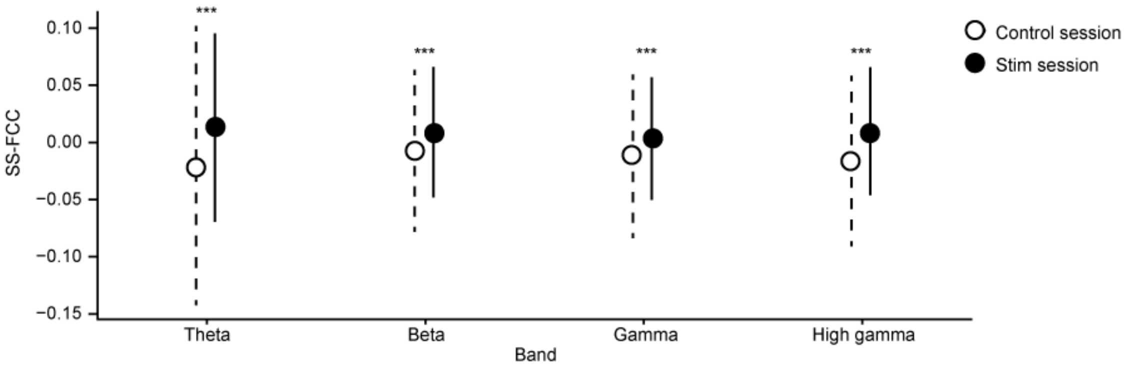


Figure S1: The average SS-FCC of stimulation experiments is higher than that of control experiments in all frequency bands. Empty circles indicate control session means while filled circles indicate stimulation session means. Dashed lines indicate standard deviation of control session means and solid lines indicate standard deviation of stimulation session means.

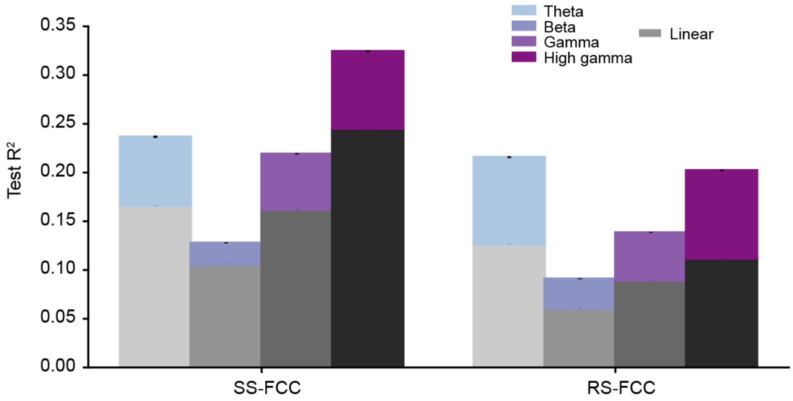


Figure S2: Accuracy of nonlinear models (colored bars) versus linear models (greyscale bars) on test data. Errors indicate standard deviation.

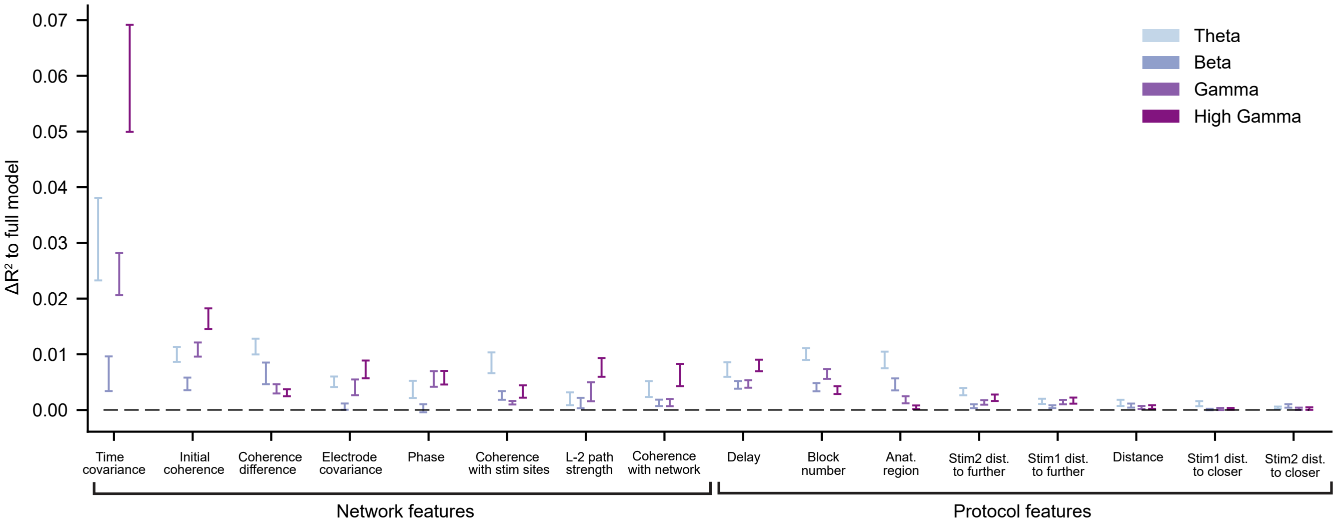


Figure S3: Feature importances for the RS-FCC models.

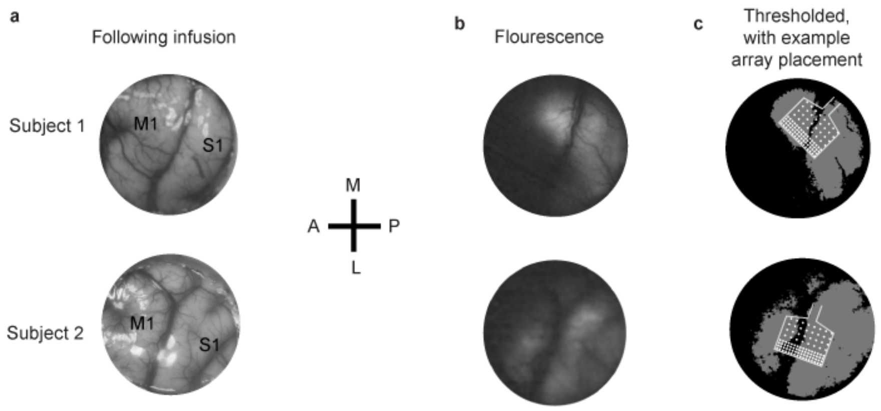


Figure S4: Widespread optogenetic expression verified through fluorescence. **a**, Images of the sensorimotor cortices of the two subjects used in our experiments. **b**, Raw images of fluorescent expression indicating optogenetic expression. **c**, Thresholded images of fluorescence shown in **b**, with schematic of recording array indicating example array placement for experimental sessions.

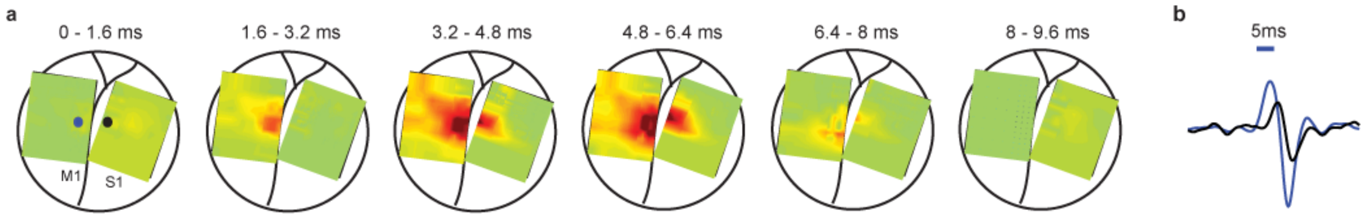


Figure S5: Spatiotemporal propagation of neural activity elicited by optogenetic activation. **a**, Heatmaps of neural activity on the cortical surface in response to optogenetic stimulation. For these data, two recording arrays were used instead of one, in order to capture a larger spatial scale of activity propagation. One µECoG array was placed on S1 and one was placed on M1, and optogenetic stimulation was delivered to an M1 site indicated by the blue circle. The ensuing neural response was filtered to the High-gamma band in order to show local neural activity. The spatiotemporal evolution of the neural activity is shown from 0 to 9.6 ms. **b**, LFP traces in the High-gamma band evoked by the stimulation as measured at the M1 stimulation site (blue) and a non-stimulated S1 site (black), corresponding to the blue and black circles in **a**.

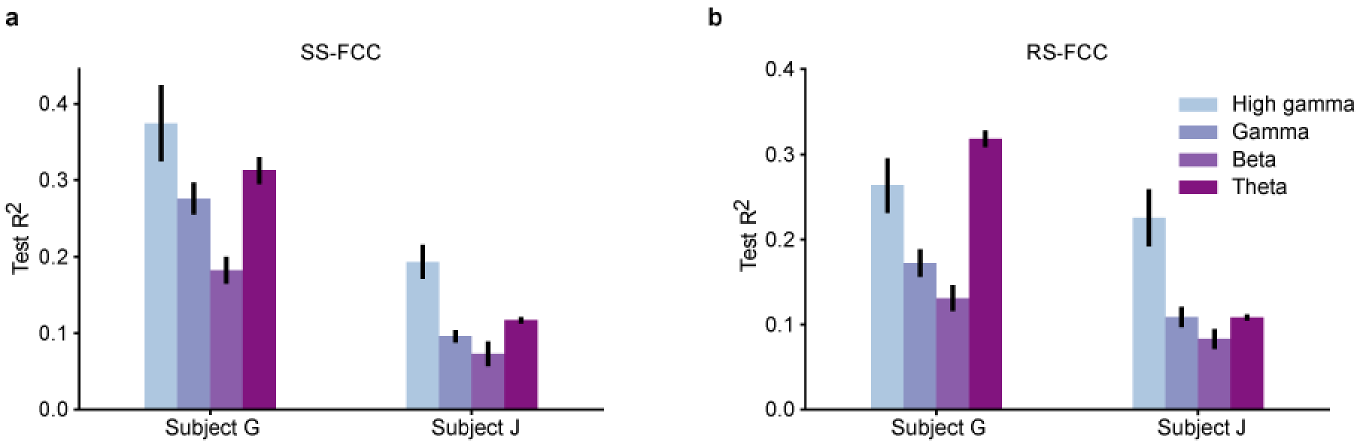


Figure S6: Accuracy of subject specific models for (**a**) SS-FCC and (**b**) RS-FCC. Data from single subjects were split into train and test data and the test accuracy is shown. Error bars indicate standard deviation.
